# RAS-inhibiting biologics identify and probe druggable pockets including an SII-α3 allosteric site

**DOI:** 10.1101/2020.06.04.133728

**Authors:** Katarzyna Z Haza, Heather L Martin, Ajinkya Rao, Amy L Turner, Sophie E Saunders, Britta Petersen, Christian Tiede, Kevin Tipping, Anna A Tang, Modupe Ajayi, Thomas Taylor, Keri M Fishwick, Thomas L Adams, Thembaninkosi G. Gaule, Chi H Trinh, Matthew Johnson, Alexander L Breeze, Thomas A Edwards, Michael J McPherson, Darren C Tomlinson

## Abstract

RAS mutations are the most common oncogenic drivers across human cancers, but there remains a paucity of clinically-validated pharmacological inhibitors of RAS, as druggable pockets have proven difficult to identify. We have identified two RAS-binding Affimer proteins, K3 and K6, that inhibit nucleotide exchange and downstream signalling pathways with distinct isoform and mutant profiles. Affimer K6 is the first biologic to bind in the SI/SII pocket, whilst Affimer K3 is the first non-covalent inhibitor of the SII region, revealing a novel RAS conformer with a large, druggable SII/α3 pocket. This work demonstrates the potential of using biologics with small interface surfaces to select novel druggable conformations in conjunction with pharmacophore identification for hard-to-drug proteins.

## INTRODUCTION

The RAS family of small GTPases consists of four members, KRAS4A, KRAS4B, HRAS and NRAS, which act as bi-directional molecular switches that cycle between an inactive GDP-bound form and an active GTP-bound form^1^. Mutations in RAS are the most common oncogenic drivers, with KRAS being the most frequently affected member especially in pancreatic, lung and colon cancer^1^. This makes RAS a strong therapeutic target, but despite having been identified as a drug target for over 30 years, only recently have compounds been developed that show promise in pre-clinical trials^2^. This paucity of agents has been driven by the lack of clearly druggable pockets on the surface of RAS. However, recent work has identified two pockets that may be amenable for drug binding^3-11^. The first of these, the SI/II-pocket, exists between the Switch I and Switch II regions of RAS in an area involved in the binding of the nucleotide exchange factor, Son of Sevenless (SOS). Several groups have independently developed compounds that bind this pocket with varying affinities and efficacies, predominately in the micromolar range^5-8^, except for BI-2852 which has nanomolar binding affinity and efficacy^11^. The second, the SII-pocket, is located under the Switch II loop and was identified using a tethered warhead approach relying on the reactive nature of the cysteine in the G12C mutant^9,10^. This pocket is not fully formed in published KRAS structures in the absence of the inhibitor; however, a groove is evident in some structures and it has been identified as a potential allosteric site computationally^3,4^. Development of tethered compounds targeting this pocket has led to the only series of RAS inhibitors currently in clinical trials^12,13^. This compound series is limited to cancers harbouring G12C mutations and it would be interesting to determine whether this cryptic groove can be exploited in other RAS isoforms and mutants.

Targeting of RAS has also been explored using biologics. Antibodies and their alternatives have been developed that bind RAS with nanomolar affinities, inhibiting nucleotide exchange, interactions with RAF, and activation of downstream pathways concurrent with negative impacts on RAS-induced cell growth and transformation^14-19^. Amongst others these include scFvs, DARPins and monobodies ^14,15,17,19,20^. Although to date some of these have been used to assist the identification of small molecules^5,7,21^, none has directly probed druggable pockets on RAS, as the majority of these biologics tend to bind over large protein interfaces^14,15,17,20^ that are difficult to mimic with small molecules^22^. As some biologics form smaller interfaces^19,23,24^ there emerges the tantalising prospect that biologics could be used as tools to identify druggable pockets and novel conformers, and could also have the potential to act as pharmacophore templates for *in silico*-informed drug discovery. Here, we explore the possibility of using Affimer proteins, an established biologic with a small probe surface formed by two variable regions^24^ known to bind at protein interaction ‘hotspots’^23,25,26^, to identify and probe RAS for druggable pockets and conformers that might be amenable to small molecule inhibition. Such a direct approach utilising small probe surfaces has not previously been used with biologics and could revolutionise drug discovery, exemplifying a novel pipeline for small molecule design that has the potential to unlock the vast number of currently ‘undruggable’ proteins.

Here, we demonstrate the use of Affimer proteins to identify and directly probe two druggable pockets on wild-type KRAS associated with inhibition of nucleotide exchange and effector molecule binding. The Affimer that binds to the SI/SII pocket actually mimics the current small molecule inhibitors that target the pocket, providing a proof-of-principle for using Affimer-target interfaces as pharmacophore templates. The Affimer that binds the SII region selects a novel conformer of the pocket present in wild-type KRAS. The Switch II region adopts a more open position, demonstrating that selecting and targeting this site via non-covalent binding is possible. Our work demonstrates two important concepts in the use of biologics: first, biologics can act as pharmacophore templates for development of small molecule inhibitors; second, they can be used to select for, and stabilise, conformations that are only present as a small fraction of the conformations of the target protein in solution, particularly those that may not be present in extant crystal structures. This approach is likely to be applicable to other important therapeutic targets, and exemplifies a novel pipeline for drug discovery.

## RESULTS

### Identification and biochemical characterization of anti-RAS Affimers

Seven unique Affimer proteins that bind wild-type KRAS in both the inactive GDP-bound form and the active form, bound to the non-hydrolysable GTP analogue - GppNHp, were isolated by phage display^27^ (Supplementary Table 1). To identify inhibitors of RAS, these Affimer proteins were screened for their ability to inhibit SOS1-mediated nucleotide exchange, the primary process in RAS activation. Three of the Affimers, K3, K6 and K37, showed clear inhibition of this process (Fig. 1a), the remaining four Affimers that bound to KRAS showed partial (Affimers K19 and K68) or no inhibition (Affimers K69 and K91) of nucleotide exchange. The latter four Affimers are not discussed further. Affimer K3 displayed the greatest inhibition of nucleotide exchange on wild-type KRAS with an IC_50_ of 144 ± 94 nM, with Affimers K6 and K37 also displaying strong inhibition with IC_50s_ of 592 ± 271 nM and 697 ± 158 nM, respectively (Supplementary Table 2). Next, the abilities of the inhibitory RAS-binding Affimers, K3, K6 and K37 to disrupt the interaction of RAS with its effector protein RAF were determined by KRAS:RAF immunoprecipitation experiments (Fig. 1b and c). Again, all three Affimer proteins caused a significant reduction in the amount of KRAS immunoprecipitated, with K3 being the most potent with a 79% reduction compared to a control Affimer, while K6 and K37 showed 40% reductions (p<0.01, p<0.05 and p<0.05 respectively; One-way ANOVA with Dunnett’s post hoc test).

**Figure 1:**
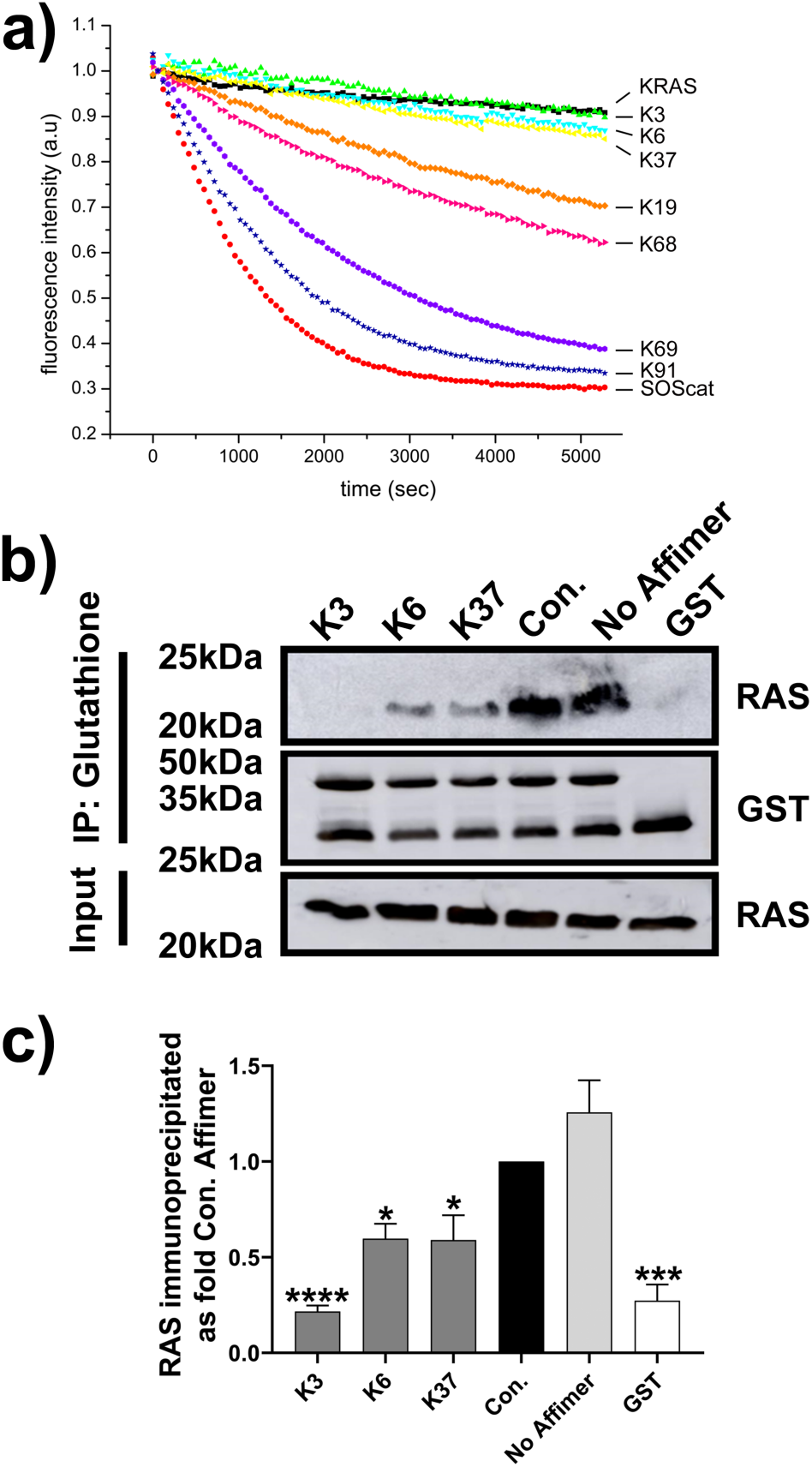
Biochemical analysis of RAS-binding Affimers. **a)** Screening of seven unique RAS-binding Affimers in a nucleotide exchange assay shows 3 Affimers, K3 (green triangles), K6 (turquoise triangles) and K37 (yellow triangles), inhibit SOS1-mediated nucleotide exchange, whilst K19 (orange diamonds) and K68 (magenta triangles) show inhibition of intrinsic nucleotide exchange and Affimers K69 (purple hexagons) and K91 (navy stars) do not inhibit nucleotide exchange. KRAS alone is shown as black squares and in the presence of SOS1^cat^ as red circles. **b)** Immunoprecipitation of KRAS with GST-RAF-RBD is inhibited by the RAS-binding Affimers, K3, K6 and K37, compared to control Affimer which does not differ from the no Affimer (PBS). GST alone does not pull-down RAS. A representative blot is shown. **c)** Quantification of b) using ImageQuantTL. Data is mean ± SEM, n = 3 independent experiments ANOVA with Dunnett’s post-hoc test * *p* < 0.05, *** *p* < 0.001, **** *p* < 0.0001. Con. – Control Affimer, RBD-RAS Binding Domain, IP – immunoprecipitation.

Having been selected against KRAS, next the specificity of the Affimer proteins for distinct RAS isoforms was assessed by nucleotide exchange assays. Whilst K6 and K37 showed no isoform specificity (Supplementary Table 2), Affimer K3 demonstrated a degree of isoform specificity, with weaker inhibition of HRAS and no measurable inhibition of NRAS with IC_50_ values of 144 ± 94 nM for KRAS, 2585 ± 335 nM for HRAS and not obtainable for NRAS, respectively. The effects of the Affimer proteins on mutant forms of KRAS were also evaluated by nucleotide exchange assays with recombinant G12D, G12V and Q61H KRAS mutants. Only Affimer K3 displayed a distinct mutant profile with 20-fold weaker inhibition of Q61H (IC_50_ = 3005 ± 865 nM), suggesting specificity towards wild-type KRAS and G12 mutations.

### Affimer proteins bind to intracellular RAS and inhibit downstream signalling

We then examined whether the Affimer proteins retained their ability to interact with and inhibit RAS in human cells by using transiently transfected HEK293 cells and His-tagged Affimer proteins. Affimers K3, K6, and K37 all showed the ability to pull down endogenous RAS, while the control Affimer showed no such activity (Fig. 2a), demonstrating binding within live cells. To understand the effects of Affimer protein binding to endogenous RAS on downstream signalling, activation of the MAPK pathway was explored. HEK293 cells were transiently co-transfected with plasmids encoding turbo-GFP (tGFP)-tagged Affimers and FLAG-tagged ERK1 expressing constructs and were then stimulated with epidermal growth factor (EGF) – this stimulation normally induces phosphorylation of ERK1 via MAPK. All three Affimer proteins significantly reduced phosphorylation of the recombinant ERK1 (One-way ANOVA with Dunnett’s post-hoc test *p*<0.001 K3, *p*<0.0001 K6 and *p*<0.0001 K37, respectively). However, the effect of Affimer K3 was lower in magnitude than those of K6 and K37, with K3 showing only a 31% reduction compared with an 85% reduction for K6, and a 69% reduction for K37 (Fig. 2b and c).

**Figure 2:**
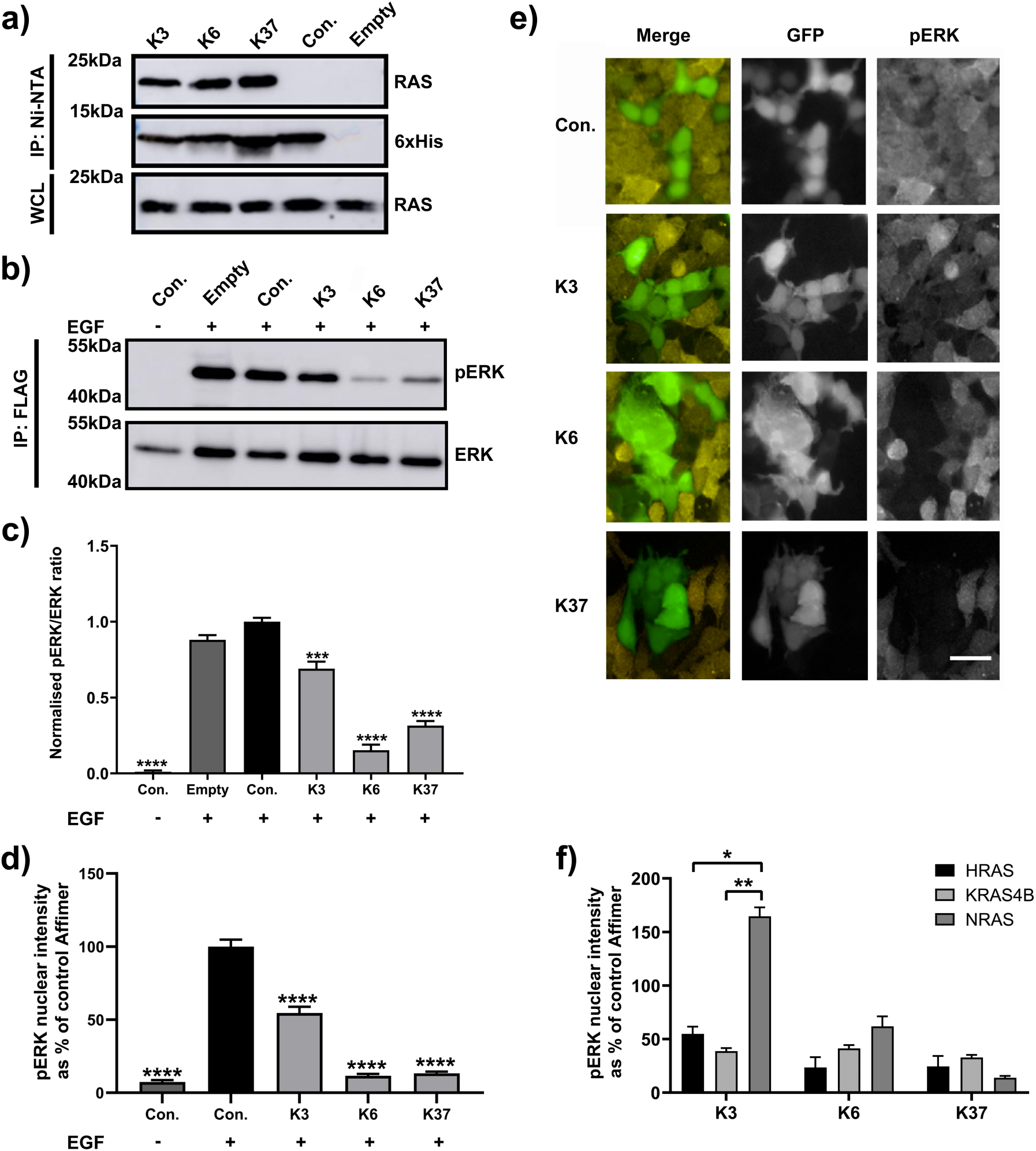
Affimers bind to intracellular RAS and inhibit downstream signalling. **a)** Immunoprecipitation of endogenous RAS from HEK293 cells transiently expressing Affimers intracellularly. RAS-binding Affimers, K3, K6 and K37, pulled down endogenously expressed RAS, whilst the control Affimer did not. **b)** HEK293 cells were co-transfected with FLAG-ERK1 plasmid and pCMV6 encoding tGFP tagged Affimers. Twenty-four hours post transfection, cells were serum-starved for 1h, then treated with 25 ng/ml EGF for 5 min. FLAG-ERK1 was precipitated from cell lysates using anti-FLAG beads and analysed for phosphorylation by immunoblotting with anti-ERK and anti-phospho-ERK antibodies. **c)** Quantification of results in b showing that Affimers K6 and K37 reduced ERK phosphorylation by over 60% while Affimer K3 reduced it by 30%. **d)** RAS-binding Affimers reduce EGF-induced phosphorylation and nuclear translocation of endogenous ERK in HEK293 cells as measured by immunofluorescence as a percentage of the control Affimer, with Affimers K6 and K37 showing inhibition of over 80% whilst Affimer K3 inhibits by 50%. **e)**. Representative images of the effects of RAS-binding Affimers, K3, K6 and K37, and the control Affimer on EGF-stimulated upregulation of pERK in HEK293 cells. **f)** RAS-binding Affimers inhibition of EGF-induced phosphorylation and nuclear translocation of endogenous ERK in mouse embryonic fibroblasts (MEFs) expressing single human RAS isoforms as a percentage of the control Affimer. Affimers K6 and K37 shown inhibition in all RAS isoforms, whilst Affimer K3 inhibited KRAS and HRAS to lesser degree with no inhibition of NRAS. Scale bars are 50µm. Data are mean ± SEM, n = 3 independent experiments, ANOVA with Dunnett’s post-hoc test * *p* < 0.05 ** p < 0.01, *** *p* < 0.001, **** *p* < 0.0001. Con. – Control Affimer, EGF – epidermal growth factor, tGFP – turbo green fluorescent protein, WCL – whole cell lysate, IP – immunoprecipitation, Empty – transfection reagents only.

To further study the impacts of RAS-binding Affimer proteins on ERK phosphorylation, we developed an immunofluorescence assay to allow the phosphorylation levels of the endogenous ERK to be examined. HEK293 cells were transiently transfected with tGFP-tagged Affimer-expressing constructs, stimulated with EGF, fixed and stained with an anti-phospho-ERK (pERK) antibody and analysed for alterations to nuclear pERK levels. In accordance with results from the immunoprecipitation experiments, expression of all three Affimer proteins resulted in significant reductions in pERK levels (One-way ANOVA with Dunnett’s post-hoc test p<0.001 for all three Affimers), with K3 having a less pronounced effect (ca. 50% reduction for K3, compared with K6, 90% and K37, 85% reductions) (Fig. 2d and e). We speculated that the lower level of inhibition observed in K3-transfected cells was due to the RAS variant specificity of this Affimer: HEK293 cells express all RAS genes, but predominantly HRAS^28^, against which K3 is 20-fold less active. To test this hypothesis, the Affimer proteins were transfected into Mouse Embryonic Fibroblasts (MEFs) that have been engineered to express single human RAS genes (gift from William Burgen at Fredrick National Laboratory for Cancer Research, Maryland, USA). All three Affimer proteins showed robust inhibition of ERK phosphorylation in KRAS-expressing MEFs with K6 and K37 showing a similar level of inhibition in MEFs expressing HRAS and NRAS. By contrast, K3 showed variant specificity, with inhibition of NRAS significantly ablated whilst the degree of inhibition for HRAS was reduced, but not significantly when compared with that of KRAS (Two-way ANOVA with Tukey’s post hoc test *p<*0.05 for NRAS vs. HRAS and *p<*0.01 for NRAS vs. KRAS) (Fig. 2f). Thus, the cellular data support the biochemical data that Affimer K3 has a preference for KRAS over NRAS, with HRAS values being intermediate.

To evaluate the impact of the Affimer proteins on MAPK signalling in the presence of mutant, oncogenic forms of KRAS, the following cancer cell lines were utilised: Panc 10.05 (KRAS G12D), SW620 (KRAS G12V) and NCI-H460 (KRAS Q61H). As anticipated from the biochemical data, all three Affimer proteins showed robust inhibition of FLAG-ERK1 phosphorylation in Panc 10.05 and SW620 cells (*p*<0.0001 and *p*<0.001 respectively One-way ANOVA with Dunnett’s post-hoc test) (Fig. 3a and b). However, in the Q61H mutant NCI-H460 cells despite all three Affimers showing inhibition of FLAG-ERK1 phosphorylation, the magnitude was reduced by 30-40% for K3 and K37 compared to the other mutant cell lines (p<0.05 One-way ANOVA with Dunnett’s post hoc test) (Fig. 3c) supporting the biochemical data that the Affimer proteins show mutant specificities. The impacts of the Affimer proteins on mutant KRAS were also tested using an immunofluorescence assay in conjunction with MEF cells expressing KRAS mutants; G12D, G12V and Q61R. Only Affimer K3 in the Q61R background showed a significant lack of inhibition of pERK nuclear intensity (p<0.05 Two-way ANOVA with Tukey’s post-hoc test) (Fig 3d), supporting the observation that Affimer K3 delineates between KRAS Q61 mutants and other KRAS variants. Together, these cellular data confirm that RAS-binding Affimer proteins are functional in cells, reducing ERK phosphorylation, and that Affimer K3 demonstrates RAS variant and mutant specificity in cells as well as *in vitro*. Given the similarities in biochemical profile and cellular activities between K6 and K37, together with a similar amino acid sequence motif (Supplementary Table 1), we postulate that they are likely to bind the same epitope although K37 showed less potency in the nucleotide exchange assays. We therefore focused on Affimer proteins K3 and K6 in our subsequent structural studies to interrogate the interactions of Affimers K3 and K6 with RAS.

**Figure 3:**
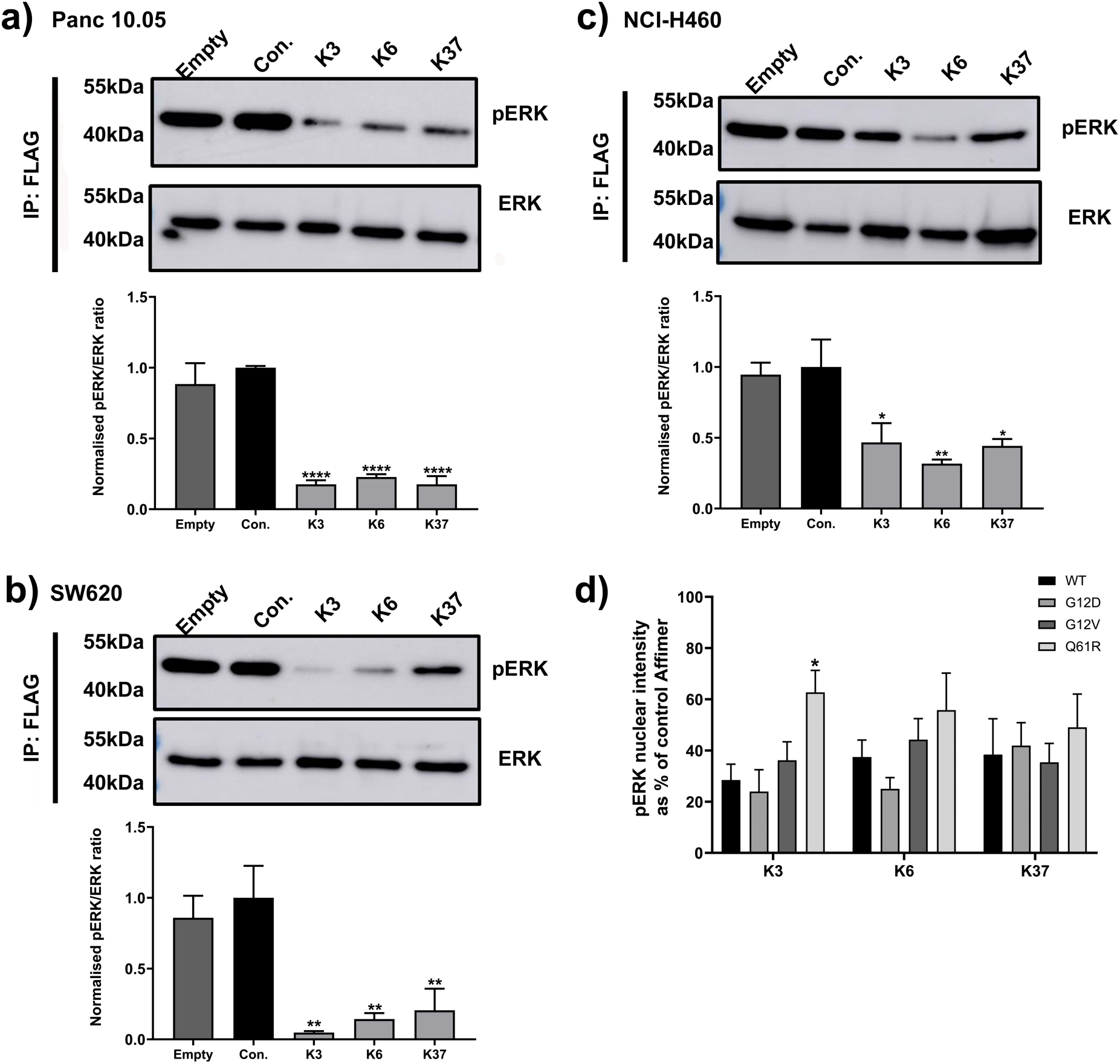
RAS-binding Affimers show different mutant specificities. **a)** Panc 10.05 (KRAS G12D), **b)** SW620 (KRAS G12V) and **c)** NCI-H460 (KRAS Q61H) cells were co-transfected with FLAG-ERK1 plasmid and pCMV6 encoding tGFP tagged Affimers. Twenty-four hours post transfection cells were serum-starved for 1h. FLAG-ERK1 was precipitated from cell lysates using anti-FLAG beads and analysed for phosphorylation by immunoblotting with anti-ERK and anti-phospho-ERK antibodies. Representative blots are shown together with quantification graphs. All three RAS-binding Affimers, K3, K6 and K37, inhibit ERK phosphorylation in Panc 10.05 cells (**a)**) and SW620 cells (**b)**). The magnitude of inhibition by Affimers K3 and K37 is reduced in NCI-H460 cells (**c)**). **d)** RAS-binding Affimers inhibit EGF-induced phosphorylation and nuclear translocation of endogenous ERK in mouse embryonic fibroblasts (MEFs) expressing single human KRAS mutants (G12D, G12V, Q61R) as measured by immunofluorescence as a percentage of the control Affimer. Only Affimer K3 shows weaker inhibition in the Q61R expressing cell line. Data are mean ± SEM, n = 3 independent experiments. One-way ANOVA with Dunnett’s post-hoc test (a-c), two-way ANOVA with Tukey’s post-hoc (d), * *p* < 0.05 ** p < 0.01, *** *p* < 0.001, **** *p* < 0.0001. Con. – Control Affimer, EGF – epidermal growth factor, tGFP – turbo green fluorescent protein, IP – immunoprecipitation, Empty – transfection reagents only, WT – wild type.

### K6 binds the SI/II hydrophobic pocket on KRAS

The crystal structure of Affimer K6 in complex with GDP-bound wild-type KRAS was determined at 1.9 Å resolution, revealing that Affimer K6 binds to a shallow hydrophobic pocket on KRAS between the switch regions (Fig. 4a and Supplementary Table 3). The Affimer K6 binding sites overlaps that of SOS1 providing structural evidence that K6 acts as a SOS1 competitive inhibitor (Supplementary Fig. 1a). The binding site further overlaps with that of the RAS-binding domain (RBD) of RAS-bound RAF (Supplementary Fig. 1b) supporting the RAS:RAF immunoprecipitation results. Affimer residues 40-45 from variable region 1 are important in the binding interface between KRAS and Affimer K6 (Fig. 4b and c). The Affimer tripeptide motif formed of P42, W43, and F44 binds the shallow hydrophobic pocket of KRAS (Fig. 4b – top left panel). The P42 residue forms no interactions with KRAS, however it is critical for function suggesting it has a geometric and structural role facilitating the interactions formed by W43 and F44. W43 of Affimer K6, and V7/L56 from KRAS form a hydrophobic cluster strengthened by Affimer residues T41 and Q45 forming hydrogen bonds with KRAS switch region residues D38/S39 and Y71, respectively (Fig. 4b -top right panel). The importance of these amino acid residues for K6 function was confirmed by mutational analysis. Individual replacement of P42, W43, F44 or Q45 by alanine reduced Affimer-mediated inhibition of nucleotide exchange (Fig. 4d). These data also revealed the importance of residues F40, N47 and R73 for the inhibitory function of Affimer K6. Indeed, complete removal of the second variable region abolished the inhibitory ability of K6 (Fig. 4d ΔVR2). This effect is most likely a result of F40, N47 and R73, and indeed Q45 forming intra-Affimer hydrogen bonds that stabilise the tripeptide, P42, W43, F44 (Fig. 4b – bottom panel). These data suggest that the functional Affimer motif responsible for binding and inhibition of KRAS is small, in agreement with the total interacting surface interface estimated by PISA analysis ^29^ of 478.3 Å^2^ for the K6:KRAS complex, a substantially smaller area than most common protein-protein interaction surfaces. The combination of a functional motif and small interaction interface provides a good basis from which to consider the development of small molecule inhibitors.

**Figure 4.**
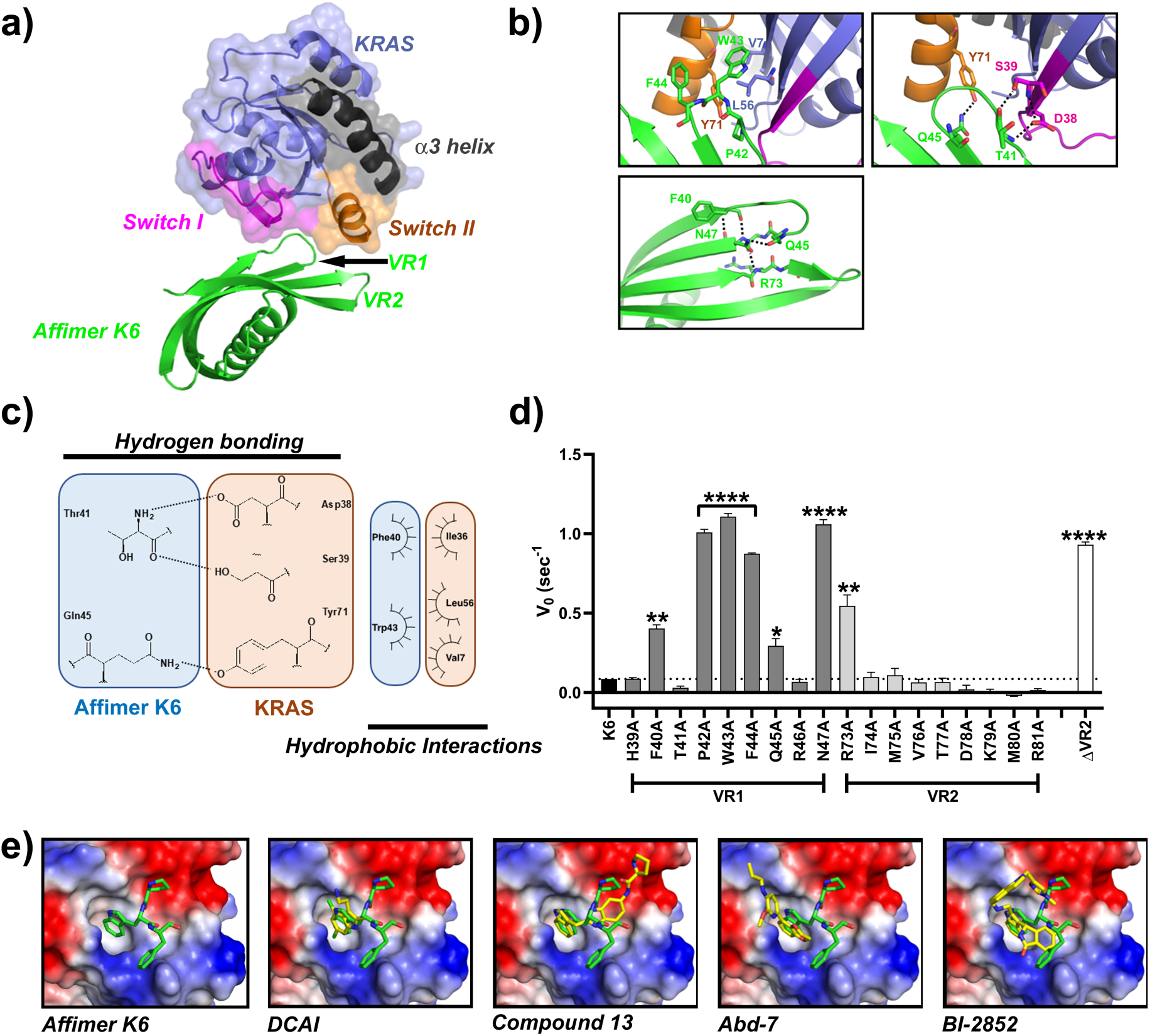
Variable region 1 of Affimer K6 binds between switch I and switch II of KRAS. **a)** Affimer K6 (green) was co-crystallized with KRASGDP (slate) and solved to a resolution of 1.9Å. The switch I (magenta), switch II (orange) and α3 helix (black) are depicted, showing their relative positioning around variable region 1 of Affimer K6. **b)** Intra- and inter-molecular interactions in the KRAS:Affimer K6 co-crystal structure are depicted; black dotted lines represent the hydrogen bonds that stabilize the critical hydrophobic contacts. **c)** Affimer K6 (VR1) and KRAS interactions shown in planar form. Hydrogen bonds are shown as black, dotted lines between the contributing atoms; additional hydrophobic interactions are represented by arcs, their spokes radiating towards the residues they contact. **d)** Alanine scanning data of the variable regions of Affimer K6 highlights Affimer residues important for inhibition of nucleotide exchange and the importance of VR2. Unaltered K6 is shown in black, variable region 1 residues are shown in dark grey, variable region 2 residues in light grey and removal of variable region 2 (ΔVR2) in white. **e)** Comparison of Affimer K6 tripeptide, P42, W43, F44, (green) with the small molecules (yellow) that bind the same SI/SII pocket. Data is mean ± SEM, n=3 independent experiments, One-way ANOVA with Dunnett’s post hoc test * *p* < 0.05 ** *p* < 0.01, **** *p* < 0.0001. Images were generated in MacPyMOL v1.7.2.3, and ChemDraw Prime 16.0. VR – variable region.

Affimer K6 binds the SI/SII pocket that has been previously documented^3-8,11^, and alanine scanning has suggested a functional pharmacophore from K6 is responsible for the observed inhibition. This PWFQxN peptide motif is also present in Affimer K37 and it is thus likely that K37 interacts in a similar manner to K6, but this remains to be confirmed. This binding pocket had previously been identified, and a number of small molecules exist to target it, including DCAI^6^, compound 13^8^, Abd-7^7^ and BI-2852^11^. We measured the affinity of K6 for both GDP- and GppNHp-bound KRAS by SPR to test whether this was comparable to these small molecules. Affimer K6 bound both forms of KRAS with low nanomolar affinities and showed a significant preference for GDP-bound KRAS (K_D_ =1.36 ± 0.87 nM for GDP and K_D_ = 7.88 ± 1.09 nM for GppNHp, Student T-test *p=*0.0095). Thus Affimer K6 has a 10-fold higher affinity for KRAS than Abd-7^7^, the strongest-binding small molecule with a K_D_ of 51 nM (c.f 750 nM for BI-2852^11^ and 1.1 mM for DCAI^6^ and 340 µM for compound 13^8^). We also inspected the SI/SII-binding small molecules for structural similarities to the K6 pharmacophore (Fig. 4e). All of the small molecules have an aromatic ring that inserts into the pocket and which is reproduced by the side chain of W43 of Affimer K6. However, only K6 and BI-2852 appear to interact across the whole of the pocket surface. This suggests that additional points of interaction may underlie efficacy, as BI-2852 is the most potent of the compounds to date^6-8,11^ and Affimer K6 shows a similar degree of potency to BI-2852, both showing IC_50_ values in the nanomolar range for inhibition of nucleotide exchange (IC_50_ = 592 ± 271 nM for Affimer K6 and IC_50_ = 490 nM for BI-2852^11^).

### K3 locks KRAS in a novel conformation to reveal a SII/α3 pocket

Affimer K3 lacks the PWFQxN motif of Affimers K6 and K37, and has a different biochemical profile in terms of mutant and isoform specificities. The underlying reasons for this were confirmed by determining the crystal structure of Affimer K3 in complex with the GDP-bound form of KRAS to 2.1 Å resolution (Fig. 5a and Supplementary Table 3). Affimer K3 binds KRAS with high affinity irrespective of the nucleotide bound (K_D_ =59.4 ± 15 nM for GDP-bound and K_D_ = 44.4 ± 0.8 nM for GppNHp-bound). Variable region 1 binds KRAS between SII and the α3 helix, with residues 41-46 being crucial for interaction (Fig. 5b and c) and inhibition. Indeed, the functional importance of these Affimer residues was highlighted by mutational analysis, as individual alanine substitutions abolished inhibition of SOS1-mediated nucleotide exchange (Fig. 5d). Affimer K3 residue D42 bridges the gap between SII and α3 helix, forming hydrogen bonds with R68 and Q99, respectively (Fig 5b – top left panel). The KRAS:K3 binding is strengthened by Affimer K3 residue D46 binding both Q99 and R102 of the α3 helix (Fig 5b – top left panel), as well as Y45 of K3, forming a hydrogen bond with E62 of SII (Fig 5b – top right panel). This forms a large binding surface allowing Affimer K3 residues I41 and I43 to form hydrophobic interactions with V103 and M72, and V9 of KRAS respectively (Fig 5b – bottom left panel). In addition, the backbone carbonyl of W44 forms a hydrogen bond with H95 of the α3 helix as well as orientating its indole side chain towards KRAS so that it is packed against the residues of Q61, H95 and Y96 (Fig 5b – top right panel). The involvement of Affimer K3 residue W44 in binding H95 may explain the specificity of K3 for KRAS seen in the biochemical and cellular data, as H95 is a unique residue only present in KRAS and not in HRAS and NRAS. The importance of H95 in mediating Affimer K3 selectivity for KRAS was confirmed by immunoprecipitation assays of mutants H95Q and H95L to mimic the corresponding residues in HRAS and NRAS. Mutating H95 affected the ability of Affimer K3 to immunoprecipitate KRAS, with mutation to glutamine (as found in HRAS) reducing the amount of RAS pulled down by 45% and mutation to a hydrophobic leucine residue (as found in NRAS) abolishing RAS immunoprecipitation completely thus giving support to our previous observations (Fig. 5e).

**Figure 5.**
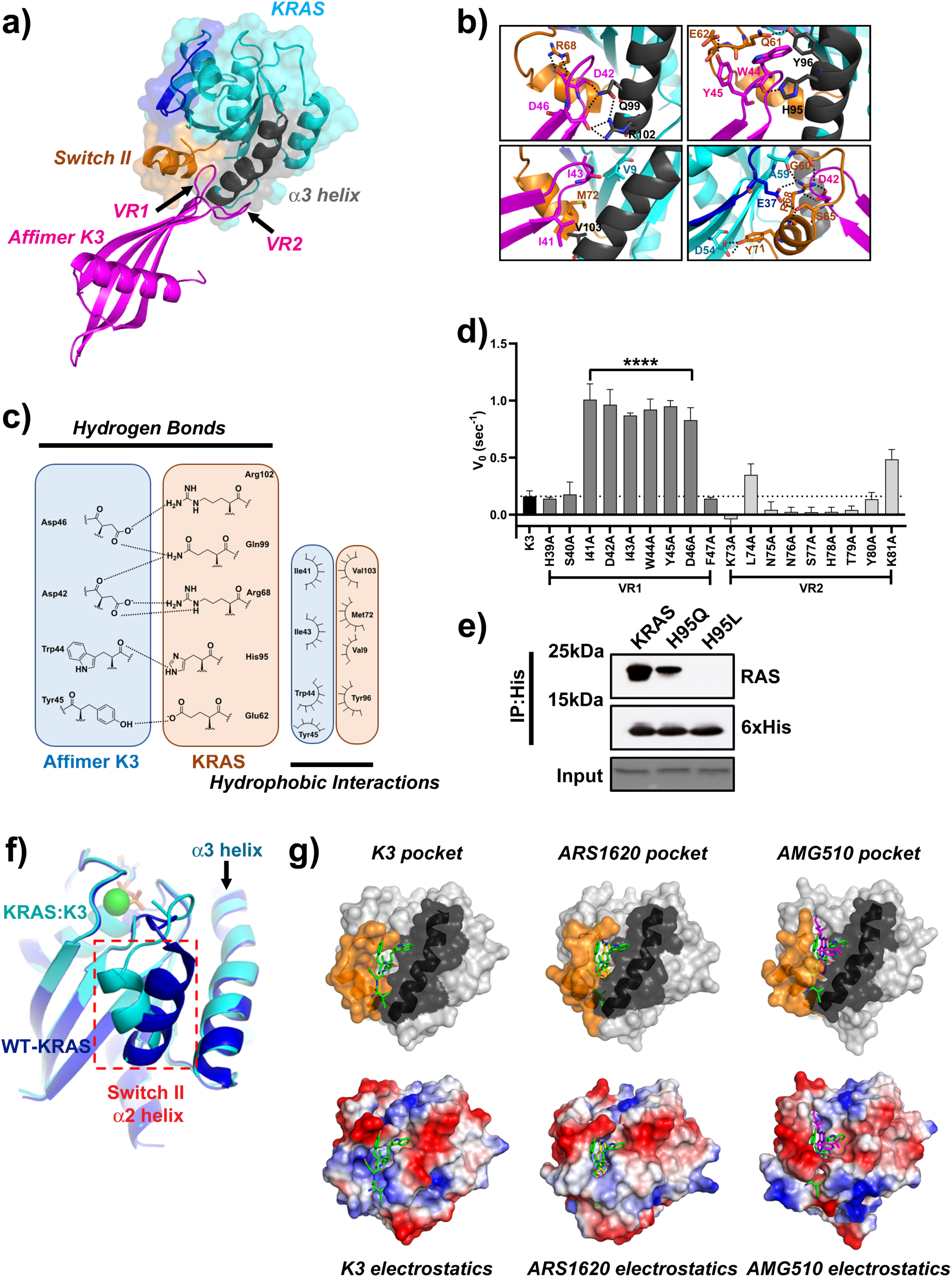
Variable region 1 of Affimer K3 explores a druggable pocket between the switch II region and α3 helix of KRAS. **a)** Affimer K3 (magenta) was co-crystallized with KRASGDP (cyan) and solved to a resolution of 2.1Å. The switch I (deep blue), switch II (orange) and α3 helix (dark grey) are depicted, showing their relative positioning around variable region 1 of Affimer K3. **b)** Intra- and inter-molecular interactions in the KRAS:Affimer K3 co-crystal structure are depicted; dotted lines (black/white) represent hydrogen bonds. **c)** All intermolecular interactions are shown in planar form. Hydrogen bonds are shown as black, dotted lines between the contributing atoms; additional hydrophobic interactions are represented by arcs, their spokes radiating towards the residues they contact. **d)** Alanine scanning data of the variable regions of Affimer K3 highlights Affimer residues important for inhibition of nucleotide exchange. Unaltered K3 is shown in black, variable region 1 residues are shown in dark grey and variable region 2 residues in light grey. **e)** Mutation of KRAS H95 affects the ability of Affimer K3 to bind, H95Q and H95L represent the residues in HRAS and NRAS respectively, a representative blot is shown. **f)** Binding of Affimer K3 causes a conformational shift to Switch II compared to WT-KRAS^GDP^. The KRAS molecule (cyan) from KRAS:Affimer K3 co-crystal structure was overlaid with WT-KRAS^GDP^ (deep blue; PDB code: 4OBE). Conformational shifts were observed in the switch II region (red-dotted box). **g)** Alterations in the conformation of the Switch II region (orange and α3 helix (black) (top row) and the corresponding alterations in the electrostatics (bottom row). Residues 41-45 of Affimer K3 (green) shown with KRASGDP (left-hand panels), overlaid with the co-crystallized KRAS:ARS1620 structure (middle panels, PBD: 5V9U) and KRAS:AMG510 structure (right-hand panels, PDB: 6OIM) (ARS1620 is shown in yellow and AMG510 is shown in magenta). Data is mean ± SEM, n=3 independent experiments, One-way ANOVA with Dunnett’s post hoc test **** *p* < 0.0001. Images were generated in MacPyMOL v1.7.2.3, and ChemDraw Prime 16.0. VR – variable region.

The inter-molecular interactions described above cause significant conformational shifts in the effector lobe of KRAS, most notably, the α2 helix of SII being forced further from the α3 helix, as compared to the WT-KRAS^GDP^ crystal structure (PDB code: 4OBE) (Fig 5f). The observed conformational shift originates from the flexible glycine residue (G60) of the DxxGQ motif of SII. The N-terminal loop of SII is further rearranged, orienting itself over the K3 binding motif towards the α3 helix. These changes in SII conformation not only generate a larger binding surface, but also facilitate a number of intra-molecular hydrogen bonds in KRAS, distinct to WT-KRAS^GDP^ (PDB code: 4OBE). Affimer K3 induces conformational changes such that the α2 helix Y71 side chain flips to interact with D54 on the β3 strand. Furthermore, we postulate that the binding of Affimer K3 residue D42 to KRAS R68, by a salt bridge interaction, shifts the R68 residue into an orientation necessary to facilitate a hydrogen bonding network between E37 of SI, A59, G60, S65 and R68 of SII thereby (Fig 5b – bottom right panel), stapling the SII region to the SI site. This hydrogen bonding network is not seen in WT-KRAS^GDP^ (PDB Code: 4OBE), or WT-KRAS^GppNHp^ (PDB code: 6GOD). At the RAS:RAF-RBD interface (PDB code: 4G0N), E37 acts as the propagating residue responsible for a hydrogen bonding/salt bridge network, terminating at the RAF-RBD residue R100^30^. It is possible that the positioning of KRAS residue E37 whilst interacting with S65 and R68 is now unable to interact with RAF-RBD and thus may explain the ability of K3 to inhibit the KRAS:RAF interaction as seen in the immunoprecipitation experiments. In addition, the conformational shift of SII upon K3 binding has orientated the M67 side chain of α3 helix towards the KRAS:RAF interface that could give rise to steric clashes with RAF-RBD residues such as R67 (Supplementary Fig 1d). Furthermore, as SII acts as a main anchor point for SOS1 binding,^31^ the significantly reduced flexibility of this site may, together with occlusion of the Cdc25 domain of SOS1 forming a steric clash (Supplementary Fig 1c), underlie the K3-mediated inhibition of SOS1-mediated nucleotide exchange. The intra-molecular hydrogen bonds between R68/E37, and Y71/D54 are also seen in the inactive WT-HRAS^GDP^ structure (PDB code: 4Q21) ^32^, suggesting K3 binding locks KRAS in an inactive conformation by stapling the switch regions together through induced hydrogen bonding, reducing conformational dynamics required for its activity.

The dynamic freedoms of SII are further reduced by the folding of the N-terminal loop of SII over the K3 binding motif resulting in a novel hydrogen bond interaction between Q61 of SII and Y96 of the α3 helix. This locks Q61 in a novel position distal to the active site. ^30,33^ This involvement of Q61 supports the biochemical and cellular data, showing the loss of inhibition of nucleotide exchange and a reduction in EGF-stimulated pERK nuclear translocation in NCI-H460 cells and Q61R MEFs, when Q61 is replaced with a histidine or arginine residue (KRAS ^Q61H/Q61R^).

Thus, Affimer K3 has identified a conformer of wild-type KRAS that generates a druggable pocket, with an estimated interface area of 790.6 Å (PISA (EMBL-EBI)^29^ analysis), buried between the SII region and the α3 helix, not previously observed in wild-type KRAS. A similar pocket has previously been documented in the KRAS G12C mutant, together with a small molecule series that inhibits RAS function via binding at this pocket^9,10,12^ (Fig. 5g). Of this series, the most recently published AMG510 compound has reached clinical development^12^. The compound is covalently tethered to the C12 residue and explores the same SII/α3 helix pocket. We hypothesise that AMG510 induces a similar mode of inhibition seen by K3, whereby the switch regions are stabilised by the ligand. However, it is clear that although the AMG510 compound and Affimer K3 explore the same cryptic groove, the conformations of SII lead to distinct pocket conformations with widely different electrostatics. Affimer K3 stabilises a more open conformation compared to the closed conformation seen with AMG510 (Fig. 5g). Further to this, binding of K3 to KRAS decreases flexibility by inducing hydrogen bond interactions between SI/SII, and SII/α3 helix, that are not present in the KRAS^G12C^:AMG510 structure (PDB code: 6OIM). The shape, size and physiochemical composition of the pocket identified by Affimer K3 suggests a potential druggable site^34,35^. The K3 data shows that we have isolated a non-covalent KRAS binder and have identified a druggable pocket and pharmacophore combination through which to inhibit KRAS preferentially over other RAS isoforms.

## Discussion

We have isolated RAS-binding Affimer reagents that inhibit RAS both *in vitro* and in cells. The Affimer proteins generated show nanomolar affinities for KRAS together with IC_50_ values in the nanomolar range for inhibition of SOS1-mediated nucleotide exchange. Furthermore, they are functional intracellularly demonstrating inhibition of the MAPK pathway as assessed by ERK phosphorylation levels. Structural analyses showed that the Affimer proteins interact with RAS within druggable pockets, notably identifying a novel pocket between the Switch II region and the α3 helix, with a non-covalent binder. Thus, we have exemplified a new site for the development of compounds to inhibit KRAS, together with a pharmacophore as a starting point for this approach.

The biochemical and cellular profiles of the Affimer proteins used in this study are comparable with biologics that have previously been identified that also inhibit RAS, again with nanomolar affinities and IC_50_ values^14,15,17-20,36^. However, the majority of these do not distinguish between RAS variants, and/or mutants, and structural analyses reveal that these pan-RAS inhibitors are binding in the Switch I/II region, except for the NS1 monobody that binds the α4-β6-α5 dimerization domain^18^, and the DARPins K13 and K19^14^ (Supplementary Fig. 2) as discussed below. The binding positions of the scFV, iDab6 and the DARPins K27 and K55 all span the SI/SII pocket^15,20^, which is the location of Affimer K6 binding; however, none of these other biologics have been shown to protrude into the pocket. Indeed, structural analysis of the K6:KRAS complex showed that a tripeptide motif, P42, W43 and F44, inserts into this SI/SII pocket in a manner that mimics the binding of known small molecules targeting this pocket. The aromatic indole ring of W43 extends into the pocket, a motif seen with all the compounds targeting this site. The interactions of the compounds are then diverse compared to Affimer K6 (Fig. 4e). It would be interesting to determine if compounds based on the K6 pharmacophore were more potent than the current compounds, as K6 binds with a higher affinity than any published reagent and shows comparable inhibition^5-8,11^.

Thus, Affimer K6 demonstrates that the use of biologics with small interaction interfaces can not only bind and inhibit difficult-to-drug proteins, but also identify and probe druggable pockets on such proteins, potentially acting as templates for small molecules. Importantly, the K3 Affimer shows inhibition of RAS, but also demonstrates a preference for KRAS over the HRAS and NRAS variants. To our knowledge, the only other biologics to express such RAS variant specificity are DARPins K13 and K19^14^. This preferential behaviour is underpinned by the involvement of the H95 residue unique to KRAS; mutation of this residue abolished binding of both Affimer K3 and the DARPins K13 and K19^14^. This ablation was more complete with mutation to glutamine for the DARPins, but was still significant with Affimer K3. These differences may in part be due to the distinct binding locations of the DARPins K13 and K19 on the allosteric lobe side of H95, whereas K3 binds on the effector lobe side and locks KRAS in a conformation where a pocket is revealed^14^. The residues involved in this pocket, specifically Q61, underlie the mutational preferences of Affimer K3 for wild-type/G12 mutants vs. Q61; this selectivity is not seen with DARPins K13 and K19^14^.

The pocket revealed by Affimer K3 binding is a previously unseen conformer of the SII pocket^3,4,9,10,12,13^, which we have termed the SII/α3 pocket, and it coincides with a cryptic groove identified computationally^4,12^. The SII pocket has previously been targeted by the covalently-tethered KRAS^G12C^ inhibitors, the ARS series, and the most recent iterations that are in clinical trials^9,10,12,13^ demonstrating the clinical importance of this pocket. Whilst this compound series has yielded the most clinically promising RAS-inhibitors to date, its dependence on covalent tethering to C12 restricts its utility to KRAS^G12C^ mutant cancers only. Affimer K3 is, to our knowledge, the only RAS inhibitor to bind to a SII-derived pocket non-covalently. K3 demonstrates a similar degree of *in vitro* potency to AMG510 with IC_50_ values for nucleotide exchange of 0.15 µM and 0.09 µM, respectively^12^, suggesting that targeting this region is a good approach for allosteric inhibition of RAS.

As Gentile *et al*^37^ noted, for binding to the SII pocket, a substituted phenolic ring is required for insertion within the subpocket formed by V9, R68, D69 and M72, and to form hydrogen bonds with R68 and D69 residues. Affimer K3 fulfils these criteria with the aromatic ring of W44 extending into this subpocket and its surrounding residues, S40 and D42 forming the necessary hydrogen bonds, thus the SII/α3 pocket shares a key subpocket with the SII pocket. Indeed, K3 also forms hydrogen bonds with H95 and Q99 in common with AMG510, albeit with different orientations of H95. Nevertheless, the positioning of the α2 helix of SII is significantly different with K3 inducing a conformation where the α2 helix distal to the α3 helix, and AMG510 a more closed conformation where the α2 helix is semi-distal to the α3 helix, but the loop region of SII is held across the pocket due to hydrogen bonding between AMG510 and KRAS E63. Whilst ARS-1620, the most potent ARS compound, leaves the helices proximal to one another as seen in WT-KRAS^GDP^ (PDB: 4OBE), (Fig.5g)^10,12^. These differences suggest that there may be an extended pocket area for small molecules based on the K3 pharmacophore, as identified by mutational analysis that is targetable to achieve the first non-covalent small molecule inhibitors of KRAS via the SII/α3 pocket.

Our work presented here demonstrates a novel concept of using biologics that bind with a relatively small interface as precursors for the development of small molecule inhibitors for difficult-to-drug proteins. The Affimer proteins identified in this study inhibited RAS, by binding to shallow pockets previously identified, or pockets derived from those previously identified, with comparable affinities and *in vitro* efficacies to the best small molecules available that target these pockets. This highlights the ability of Affimer proteins to select novel conformers of target proteins to reveal druggable regions on protein surfaces concurrent with pharmacophore identification. Indeed, it will be interesting to use the pharmacophore motifs identified in this study as templates for novel series of RAS-binding small molecules and for potential hit-to-lead optimisation using an Affimer-target NanoBRET system, as has been previously achieved with RAS-binding biologics^5,7^. The approach utilised in this study is likely to be applicable to other important therapeutic targets, and provides a novel pipeline for drug discovery.

## Acknowledgments

This work was supported by the Wellcome Trust grant number 102174/B/13/Z, Medical Research Council grant number MR/N020952/1, EPSRC grant number EP/L015005/1 and Technology Strategy Board grant number TS/M001199/1.

## Author Contributions

DCT, MJM, TAE, ALB and MJ conceived the experimental plan. KZH, HLM, AR, and ALT contributed equally to the experimental data. SES, BP, CT, KT, AAT, MA, TT, KMF, TLA, TGG and CHT conducted experimental work. All authors performed data analysis and critically reviewed and approved the manuscript.

## METHODS

### Phage display and Affimer protein production

Selection of Affimers by phage display was performed as described previously^24^ against GDP-bound KRAS with the addition of 10mM MgCl_2_ to all buffers. RAS-binding Affimers were produced in BL21 STAR™ (DE3) *E. coli* (Life Technologies, Invitrogen) and their cross-reactivity against GppNHp-bound KRAS determined by ELISA as previously described^24^.

### Protein production

The human wild-type KRAS (Isoform b), HRAS and NRAS gene sequences (residues 1–166, with an N-terminal His-tag and C-terminal BAP-tag were synthesized by GenScript (Piscataway, USA) and cloned into pET11a. RAS mutants G12D, G12V, Q61H, H95Q and H95L were produced by Quikchange™ site-directed mutagenesis using the wild type as template. RAS proteins were produced in BL21 STAR™ (DE3) *E. coli* induced with 0.5 mM IPTG and grown overnight at 20°C at 150 rpm. Cells were harvested by centrifugation at 4816 x *g* for 15 min at 4°C and resuspended in 20mM Tris, pH 7.5, 500mM NaCl, 10mM Imidazole, 5% Glycerol, supplemented with EDTA-free protease inhibitor, 0.1mg/ml lysozyme, 1% Triton X-100 and 10U/ml Benzonase nuclease. Proteins were purified from the supernatant by Ni-NTA chromatography and size exclusion chromatography into RAS buffer (20mM Tris, 100mM NaCl, 10mM MgCl_2_, 1mM DTT, 5% Glycerol, pH 7.5). GST-thr-RAF1-RBD was provided as a gift from Dominic Esposito (Addgene plasmid # 86033) and produced as previously described^38^. GST-thr-RAF1-RBD supernatants were used for RAS:RAF immunoprecipitation. Human SOS1 catalytic domains (SOS1^cat^) coding region (residues 564-1059) with an N-terminal His-tag in pET11a were produced in BL21 Star™ DE3 *E. coli* following 0.5 mM IPTG induction and grown overnight at 25°C at 150 rpm. Cells were harvested by centrifugation at 4816 x *g* for 15 min at 4°C. Cell pellets were lysed in 20 mM Tris-HCl pH 8; 300 mM NaCl; 20 mM imidazole; 5% glycerol supplemented with 1% Triton-X100, EDTA-free protease inhibitor, 0.1 mg/ml lysozyme and 10 U/ml Benzonase nuclease. The lysate was centrifuged at 12,000 x *g* for 20 min then applied to Ni-NTA resin. Proteins were eluted using 20 mM Tris-HCl pH 8; 500 mM NaCl; 300 mM imidazole; 5% glycerol and dialysed into 50 mM Tris-HCl pH 7.5; 100 mM NaCl; 1 mM DTT.

### RAS nucleotide loading

RAS was desalted into nucleotide loading buffer (25mM Tris-HCl, 50mM NaCl, 0.5mM MgCl_2_ pH 7.5) using a Zeba spin column according to the manufacturer’s instructions (ThermoFisher). MANT-GDP (mGDP, SigmaAldrich) or GppNHp (SigmaAldrich) was added in a 20 fold excess over RAS together with DTT and EDTA to a final concentration of 1mM and 5mM respectively in a volume of 130µl and incubated at 4°C for 1h. MgCl_2_ was added then in a 9 fold excess over EDTA and incubated for a further 30 min at 4°C. Loaded RAS was then desalted into nucleotide exchange buffer (20mM HEPES pH 7.5, 150mM NaCl, 10mM MgCl_2_) using a Zeba spin column. Nucleotide loading was confirmed by native mass spectrometry.

### Guanine nucleotide exchange assays

Nucleotide exchange buffer was supplemented with 0.4 mM GTP (SigmaAldrich) and 0.5 µM SOS1^cat^ for experiments involving WT RAS, or 2 µM SOS1^cat^ for experiments involving mutant RAS proteins. The Affimer proteins were then diluted with the GTP-SOS1^cat^ supplemented nucleotide exchange buffer to make 20 µM stock solutions, which were then used for a 2-fold serial dilution series using the supplemented nucleotide exchange buffer. A 1µM stock of the WT/mutant RASmGDP protein was prepared by diluting the stock RAS in nucleotide exchange buffer supplemented with 2 mM DTT. Solutions were incubated at 37°C for 10 min before the assay. The reaction was initiated by addition of Affimer/SOS1^cat^/GTP solution to RAS/DTT containing solution. Changes in fluorescence were measured by a fluorescence spectrometer (Tecan Spark) in a Corning black, flat-bottomed, non-binding 384 well plate at 440nm every minute for 90 min. The data were then normalised to initial fluorescence reading and fit to a single exponential decay using OriginPro 9.7.0 software (OriginLab, Massachusetts). The derived rates were normalised to these of RAS-SOS1 minus RAS-only samples and fit to the Hill equation (y = START + (END − START) (xn / kn + xn)) from which the IC_50_ values were calculated.

### RAS Interaction Assays

The interaction of KRAS with RAF-RBD or Affimer K3 was assessed by immunoprecipitation using a KingFisher Flex (ThermoFisher Scientific). Glutathione magnetic agarose beads were blocked overnight at 4°C with 2x blocking buffer (Sigma, B6429) then incubated with RAF-RBD-GST supernatants for 1 h at room temperature. Simultaneously 1 µg/µl of KRAS-GppNHp was incubated with 0.6 µg/µl of Affimer or PBS (no Affimer control) in a total volume of 100 µl PBS. Beads were washed 3 times with assay buffer (125mM Tris, 150mM NaCl, 5mM MgCl2, 1mM DTT, 0.01% Tween-20, pH 8.0) and KRAS:Affimer solutions added and incubated for 1 h at room temperature. Beads were washed 4 times (15 secs/wash) with assay buffer and proteins were eluted with SDS-PAGE sample buffer (200mM Tris-HCl, 8% SDS, 20% glycerol, 20% mercaptoethanol, 0.1% (w/v) bromophenol blue, pH 7). RAS:Affimer immunoprecipitation was as for RAS:RAF immunoprecipitation except His Mag Sepharose™ Ni^2+^-NTA beads (GE healthcare^®^) were used and incubated with 20 µg of Affimer K3 for 1 h with agitation. After washing 3 times the beads were mixed with 100 µl of KRAS/KRAS(H95Q)/KRAS(H95L) lysate. For elution of K3:KRAS SDS-PAGE sample buffer was supplemented with 500 mM imidazole. Proteins were analysed by immunoblot with anti-GST-HRP (1:5000, GeneTex, GTX114099) or anti-6X His tag antibody (HRP) (1:10,000, Abcam, ab1187) and KRAS+HRAS+NRAS (1:1000, Abcam, ab206969) antibodies.

### Immunoblotting

Protein samples were separated on 15% SDS-PAGE and transferred onto nitrocellulose membranes. These were blocked with 5% milk (SigmaAldrich) in TBS-0.1% Tween 20 (TBST), incubated overnight at 4°C with primary antibody as detailed, washed 3 times with TBST and incubated for 1 h at room temperature with HRP-conjugated secondary antibody (1:10,000 goat anti-rabbit HRP, Cell Signalling Technology, CST7074S) if required. Following 3 x TBS-T washes the membranes were developed with Immunoblot Forte Western HRP (Millipore), according to the manufacturer’s instructions and imaged using an Amersham™ Imager 600 (GE Healthcare). Uncropped images of all membranes used in this manuscript are shown in Supplementary Figure 3.

### Cell culture

HEK293, Panc 10.05 and NCI-H460 cells were purchased from ECACC, UK. RAS-expressing mouse embryonic fibroblasts (MEFs) were from William Burgen at Fredrick National Laboratory, Maryland, USA. SW620 cells were from Professor Mark Hull, University of Leeds, UK. HEK293 and MEF cell lines were maintained in Dulbecco’s Modified Eagle Medium (SigmaAldrich) supplemented with 10% fetal bovine serum (FBS) (Gibco), and 4 µg/mL blastocidin S (SigmaAldrich)(MEFs expressing KRAS and NRAS isoforms) or 2.5 µg/mL puromycin (SigmaAldrich) (MEFs expressing HRAS). NCI-H460 and SW620 cell lines were maintained in RPMI-1640 media (SigmaAldrich) supplemented with 10% FBS. Panc10.05 cells were maintained in RPMI-1640 supplemented with 15% FBS and 10U insulin (SigmaAldrich). All cells were maintained at 37°C in CO_2_ and were mycoplasma free.

### RAS immunoprecipitation

4 × 10^5^ HEK293 cells/well were plated in 12 well plates and incubated at 37°C, 5% CO_2_ for 24 h before transient transfection with plasmids encoding Affimer-His constructs using Lipofectamine 2000 (ThermoFisher), as per the manufacturer’s instructions. After 48 h cells were lysed in NP-40 buffer (50 mM Tris, 150 mM NaCl, 1% NP-40 (v/v), 1x Halt™ protease inhibitor cocktail (ThermoFisher), 1x phosphatase inhibitor cocktail 2 (SigmaAldrich), pH 7.5.) and cleared lysates incubated overnight at 4 °C with Ni-NTA resin. After washing, proteins were eluted in SDS sample buffer and analysed by immunoblotting with the anti-KRAS+HRAS+NRAS or anti-6X His tag-HRP antibody.

### FLAG-ERK pull-down assays

Cells were plated into 12-well plates (1× 10^5^ cells/well for HEK293 cells, 2× 10^5^ cells/well for SW620 and NCI-H460 cells and 4 × 10^5^ cells/well for Panc10.05 cells) and incubated at 37 °C, 5% CO_2_ for 24 h before transfection with a 4:1 DNA ratio of pCMV6-Affimer-tGFP and FLAG-ERK plasmids using Lipofectamine 2000 (SW620, HEK293, NCI-H460) or X-tremeGENE 9 (Roche; Panc10.05). After 24 h cells were serum starved for 1 h, HEK293 cells were then stimulated with 25 ng/ml EGF (Gibco) for 5 min (other cell lines were not stimulated). Cells were washed with ice-cold DPBS then incubated for 10 min on ice with NP-40 lysis buffer. Cleared lysates were incubated overnight at 4°C with 20 µl anti-FLAG M2 magnetic beads (SigmaAldrich). The beads were washed 3x with TBS before protein elution by incubation at 95°C for 5 min in SDS sample buffer. Levels of ERK and pERK were then analysed by immunoblotting with anti-ERK antibody (1:2000, Abcam, ab184699) and phospho-ERK antibody (1:1000, Abcam, ab76299). Densitometry analysis used ImageJ software v.1.52 (NIH, Maryland).

### pERK Immunofluorescence Assay

Cells were plated into 96 well plates (1× 10^5^ cells/ml for HEK293 cells, 2-8 × 10^4^ cells/ml for MEFs) and incubated at 37 °C, 5% CO_2_ for 24 before transfection with pCMV6-Affimer-tGFP plasmids using Lipofectamine 2000 as per the manufacturer’s instructions. After a further 24 h cells were serum starved for 1-18 h before stimulation with EGF (25ng/mL) for 5 min, rinsed with PBS and fixed in 4% paraformaldehyde (VWR) for 15 min. Cells were rinsed with PBS and permeabilised with methanol at −20°C for 10 min, before rinsing with PBS and blocking (1% milk (SigmaAldrich) in PBS) and incubating with anti-pERK antibody (1:150 Cell Signalling Technology 4370) in blocking solution for 1 h at room temperature followed by 3x PBS rinses and incubating with Hoechst 33342 (1µg/mL Molecular Probes) and anti-rabbit AlexaFluor 546 or 568 (1:1000 Molecular Probes) in blocking solution for 1 h at room temperature. Following a final set of PBS washes, plates were scanned and images collected with an Operetta HTS imaging system (PerkinElmer) or ImageXpress Pico (Molecular Devices) at 20x magnification. Images were analysed with Columbus 2.7.1 (PerkinElmer) or MetaExpress 6.7 (Molecular Devices) software.

### Crystallization, data collection, and structure determination

Purified Affimer proteins were incubated with KRAS lysates overnight at 4°C and the complexes purified by Ni-NTA affinity chromatography and size exclusion chromatography using HiPrep 16/60 Sephacryl S-100 column (GE Healthcare). The complexes were concentrated to 24 mg/ml (KRAS:K3) and 12 mg/mL (KRAS:K6) respectively in 10mM Tris-HCl pH 8.0, containing 50mM NaCl, 20mM MgCl_2_ and 0.5mM TCEP. KRAS:K3 crystals were obtained in 2M (NH_4_)_2_SO_4_, 0.2M K Na tartrate and 0.1M tri-sodium citrate pH 5.6 by sitting drop vapour diffusion. Crystals were flash-cooled in a mixture of 75% mother liquor and 25% ethylene glycol. KRAS:K6 crystals were obtained in 0.1M C_2_H_3_NaO_2_ pH 5, 25% w/v PEG 4K, 0.2M (NH_4_)_2_SO_4_, 5% MPD by hanging-drop vapour diffusion. Crystals were flash-cooled in 30% w/v PEG 4K, 0.1M C_2_H_3_NaO_2_t, pH 5, 0.2M (NH_4_)_2_SO_4_, 20mM MgCl_2_, 5% PEG 400, 5% MPD, 5% ethylene glycol and 5% glycerol. X-ray diffraction data for the KRAS:K6 complex were recorded on beamline I04-1 at the Diamond Light Source with data for KRAS:K3 being recorded on beamline ID30A-1 at the European synchrotron radiation facility, at 100 K. Data collection statistics are reported in Supplementary Tables 3. Diffraction data were processed and scaled with the Xia2 suite of programs^39^. The KRAS:Affimer structures were determined by molecular replacement with the KRAS-GDP structure (PDB 4OBE) and an Affimer structure (PDB 4N6T) excluding the variable regions as the initial search models in the program Phaser^40^. Structures were refined using REFMAC5^41^, followed by iterative cycles of manual model building using COOT^42^. Whilst the final model of the KRAS:K6 structure contain all the residues of the variable regions, the electron density maps for residues 75-80 for both the KRAS:K3 complexes in the asymmetric unit cell were highly disorder with incomplete connectivity even when contoured at low sigma level. As a result KRAS:K3 residues S77, H78 and T79 were included in the final model but as poly-alanines. Model validation was conducted using the Molprobity server^43^. Molecular graphics were generated using MacPyMOL version 1.7.2.3. Surface area calculations were performed using the PDBePISA^29^ protein– protein interaction server. The KRAS:K6 and KRAS:K3 structures have been deposited with the PDB codes 6YR8 and 6YXW respectively.

### Affimer Affinity Measurements

Affimer affinities for KRAS were determined by surface plasmon resonance (SPR) using a BIAcore 3000 (GE Healthcare Europe GmbH). Affimer proteins with a C-terminal cysteine residue were biotinylated with the biotin-maleimide (SigmaAldrich) as previously described^24^ and immobilized on to streptavidin-coated CM5 sensor chips (Biacore). Biacore experiments were performed at 25°C in HEPES buffer (20 mM HEPES, pH 7.5, 150 mM NaCl, 10 mM MgCl_2_, 0.1% Tween 20, 0.1% Triton × 100). KRAS (bound to GppNHp or GDP) was injected at 6.25, 12.5, 25, 50, 100, 200, 400, and 800 nM at a flow rate of 5 µl min^-1^, followed by 3 min stabilization and 10 min dissociation. The on- and off-rates and K_D_ parameters were obtained from a global fit to the SPR curves using a 1:1 Langmuir model, using the BIAevaluation software. Quoted K_D_ values are the mean ± SEM of three replicate measurements.

### Alanine scanning by site-directed mutagenesis

To assess the importance of each residue in the Affimer variable regions point mutations to encode sequential alanine residues were introduced by Quikchange™ site-directed mutagenesis. Reactions contained 1x KOD polymerase reaction buffer, 0.2 mM dNTP, 2 mM MgSO_4_, 0.3 µM of forward and reverse primer, 10 ng DNA template and 1 U KOD polymerase. PCR amplification consisted of 30 cycles of 20 secs at 98°C, 10 secs at 68°C and 3.5 mins at 70°C. Samples were digested with Dpn I for 1 h at 37°C and introduced by transformation into XL1-Blue super-competent cells. DNA was extracted using QIAprep Spin Miniprep Kit as per the manufacturer’s instructions and mutagenesis was confirmed by DNA sequence analysis (Genewiz).

### Construction of K6ΔVR2 mutant

To generate Affimer K6ΔVR2 mutant, the 9 residues of the K6 VR2 were replaced with AAE. Affimer K6 VR1 and control Affimer VR2 (AAE) were amplified and subjected to splice overlap extension (SOE) PCR. The spliced product was subcloned into pET11a and Affimer K6ΔVR2 produced as previously described^24^.

### Statistical Analysis

Data were analyzed in Prism v8.1.0 (GraphPad Software). Normality was tested using Shapiro Wilk test. Data presented are mean ± SEM unless otherwise stated.

### Data Availability Statement

The Affimer constructs generated during the current study are available under a standard MTA from the University of Leeds via the corresponding author (DCT). The X-Ray crystal structures generated during and analysed during the current study are available in the PDB repository (https://www.rcsb.org).

## References

1. Hobbs, G.A., Der, C.J. & Rossman, K.L. RAS isoforms and mutations in cancer at a glance. Journal of Cell Science 129, 1287 (2016).

2. O’Bryan, J.P. Pharmacological targeting of RAS: Recent success with direct inhibitors. Pharmacol Res 139, 503–511 (2019).

3. Buhrman, G. et al. Analysis of binding site hot spots on the surface of Ras GTPase. Journal of molecular biology 413, 773–789 (2011).

4. Grant, B.J. et al. Novel Allosteric Sites on Ras for Lead Generation. PLOS ONE 6, e25711 (2011).

5. Cruz-Migoni, A. et al. Structure-based development of new RAS-effector inhibitors from a combination of active and inactive RAS-binding compounds. Proc Natl Acad Sci U S A 116, 2545–2550 (2019).

6. Maurer, T. et al. Small-molecule ligands bind to a distinct pocket in Ras and inhibit SOS-mediated nucleotide exchange activity. Proc Natl Acad Sci U S A 109, 5299–304 (2012).

7. Quevedo, C.E. et al. Small molecule inhibitors of RAS-effector protein interactions derived using an intracellular antibody fragment. Nat Commun 9, 3169 (2018).

8. Sun, Q. et al. Discovery of small molecules that bind to K-Ras and inhibit Sosmediated activation. Angew Chem Int Ed Engl 51, 6140–3 (2012).

9. Ostrem, J.M., Peters, U., Sos, M.L., Wells, J.A. & Shokat, K.M. K-Ras(G12C) inhibitors allosterically control GTP affinity and effector interactions. Nature 503, 548–51 (2013).

10. Janes, M.R. et al. Targeting KRAS Mutant Cancers with a Covalent G12C-Specific Inhibitor. Cell 172, 578-589.e17 (2018).

11. Kessler, D. et al. Drugging an undruggable pocket on KRAS. Proceedings of the National Academy of Sciences 116, 15823 (2019).

12. Canon, J. et al. The clinical KRAS(G12C) inhibitor AMG 510 drives antitumour immunity. Nature 575, 217–223 (2019).

13. Hallin, J. et al. The KRAS^G12C^ Inhibitor MRTX849 Provides Insight toward Therapeutic Susceptibility of KRAS-Mutant Cancers in Mouse Models and Patients. Cancer Discovery 10, 54 (2020).

14. Bery, N. et al. KRAS-specific inhibition using a DARPin binding to a site in the allosteric lobe. Nat Commun 10, 2607 (2019).

15. Guillard, S. et al. Structural and functional characterization of a DARPin which inhibits Ras nucleotide exchange. Nat Commun 8, 16111 (2017).

16. Khan, I., Rhett, J.M. & O’Bryan, J.P. Therapeutic targeting of RAS: New hope for drugging the “undruggable”. Biochim Biophys Acta Mol Cell Res 1867, 118570 (2020).

17. Shin, S.M. et al. Antibody targeting intracellular oncogenic Ras mutants exerts anti-tumour effects after systemic administration. Nat Commun 8, 15090 (2017).

18. Spencer-Smith, R. et al. Inhibition of RAS function through targeting an allosteric regulatory site. Nat Chem Biol 13, 62–68 (2017).

19. Spencer-Smith, R. et al. Targeting the alpha4-alpha5 interface of RAS results in multiple levels of inhibition. Small GTPases 10, 378–387 (2019).

20. Tanaka, T., Williams, R.L. & Rabbitts, T.H. Tumour prevention by a single antibody domain targeting the interaction of signal transduction proteins with RAS. Embo j 26, 3250–9 (2007).

21. Bery, N. et al. BRET-based RAS biosensors that show a novel small molecule is an inhibitor of RAS-effector protein-protein interactions. Elife 7(2018).

22. Arkin, M.R., Tang, Y. & Wells, J.A. Small-molecule inhibitors of protein-protein interactions: progressing toward the reality. Chemistry & biology 21, 1102–1114 (2014).

23. Robinson, J.I. et al. Affimer proteins inhibit immune complex binding to FcgammaRIIIa with high specificity through competitive and allosteric modes of action. Proc Natl Acad Sci U S A 115, E72–e81 (2018).

24. Tiede, C. et al. Affimer proteins are versatile and renewable affinity reagents. Elife 6(2017).

25. Hughes, D.J. et al. Generation of specific inhibitors of SUMO-1- and SUMO-2/3-mediated protein-protein interactions using Affimer (Adhiron) technology. Sci Signal 10(2017).

26. Miles, J.A. et al. Selective Affimers Recognize BCL-2 Family Proteins Through Non-Canonical Structural Motifs. bioRxiv, 651364 (2019).

27. Tang, A.A., Tiede, C., Hughes, D.J., McPherson, M.J. & Tomlinson, D.C. Isolation of isoform-specific binding proteins (Affimers) by phage display using negative selection. Sci Signal 10(2017).

28. Thul, P.J. et al. A subcellular map of the human proteome. Science 356, eaal3321 (2017).

29. Krissinel, E. & Henrick, K. Inference of macromolecular assemblies from crystalline state. Journal of Molecular Biology 372, 774–797 (2007).

30. Fetics, S.K. et al. Allosteric effects of the oncogenic RasQ61L mutant on Raf-RBD.

31. Bandaru, P., Kondo, Y. & Kuriyan, J. The Interdependent Activation of Son-of-Sevenless and Ras. Cold Spring Harbor perspectives in medicine 9, a031534 (2019).

32. Simanshu, D.K., Nissley, D.V. & McCormick, F. RAS Proteins and Their Regulators in Human Disease. Cell 170, 17–33 (2017).

33. Buhrman, G., Holzapfel, G., Fetics, S. & Mattos, C. Allosteric modulation of Ras positions Q61 for a direct role in catalysis. Proceedings of the National Academy of Sciences 107, 4931 (2010).

34. Halgren, T.A. Identifying and Characterizing Binding Sites and Assessing Druggability. Journal of Chemical Information and Modeling 49, 377–389 (2009).

35. Yaxia, Y., Jianfeng, P. & Luhua, L. Binding Site Detection and Druggability Prediction of Protein Targets for Structure-Based Drug Design. Current Pharmaceutical Design 19, 2326–2333 (2013).

36. Tanaka, T. & Rabbitts, T.H. Intrabodies based on intracellular capture frameworks that bind the RAS protein with high affinity and impair oncogenic transformation. Embo j 22, 1025–35 (2003).

37. Gentile, D.R. et al. Ras Binder Induces a Modified Switch-II Pocket in GTP and GDP States. Cell chemical biology 24, 1455-1466.e14 (2017).

38. Ren, X.D., Kiosses, W.B. & Schwartz, M.A. Regulation of the small GTP-binding protein Rho by cell adhesion and the cytoskeleton. The EMBO journal 18, 578–585 (1999).

39. Winter, G., Lobley, C.M.C. & Prince, S.M. Decision making in xia2. Acta Crystallographica Section D 69, 1260–1273 (2013).

40. McCoy, A.J. et al. Phaser crystallographic software. Journal of Applied Crystallography 40, 658–674 (2007).

41. Murshudov, G.N., Vagin, A.A. & Dodson, E.J. Refinement of macromolecular structures by the maximum-likelihood method. Acta Crystallographica Section D-Structural Biology 53, 240–255 (1997).

42. Emsley, P. & Cowtan, K. Coot: model-building tools for molecular graphics. Acta Crystallographica Section D-Biological Crystallography 60, 2126–2132 (2004).

43. Chen, V.B. et al. MolProbity: all-atom structure validation for macromolecular crystallography. Acta crystallographica. Section D, Biological crystallography 66, 12–21 (2010).

