## Supplementary Data for "RAS-inhibiting biologics identify and probe druggable pockets including an SII-α3 allosteric site"

**Supplementary Figure 1. Affimer binding locations compared with SOS1 and RAF-RBD. a)** Affimer K6 (green) occludes SOS1 (orange) binding RAS (grey).**b)** Affimer K6 (green) sterically clash with RAF-RBD (blue) binding of RAS (grey) **c)** Affimer K3 (magenta) occludes SOS1 (orange) binding RAS (grey). **d)** Affimer K3 (magenta) does not sterically clash with RAF-RBD (blue) binding of RAS (grey), but a conformational change to the Switch II region is seen compared to unbound KRASGDP. Images were generated in MacPyMOLv1.7.2.3 using RAS:SOS1 PDB:1BKD and RAS:RAF-RBD PDB:4G0N.

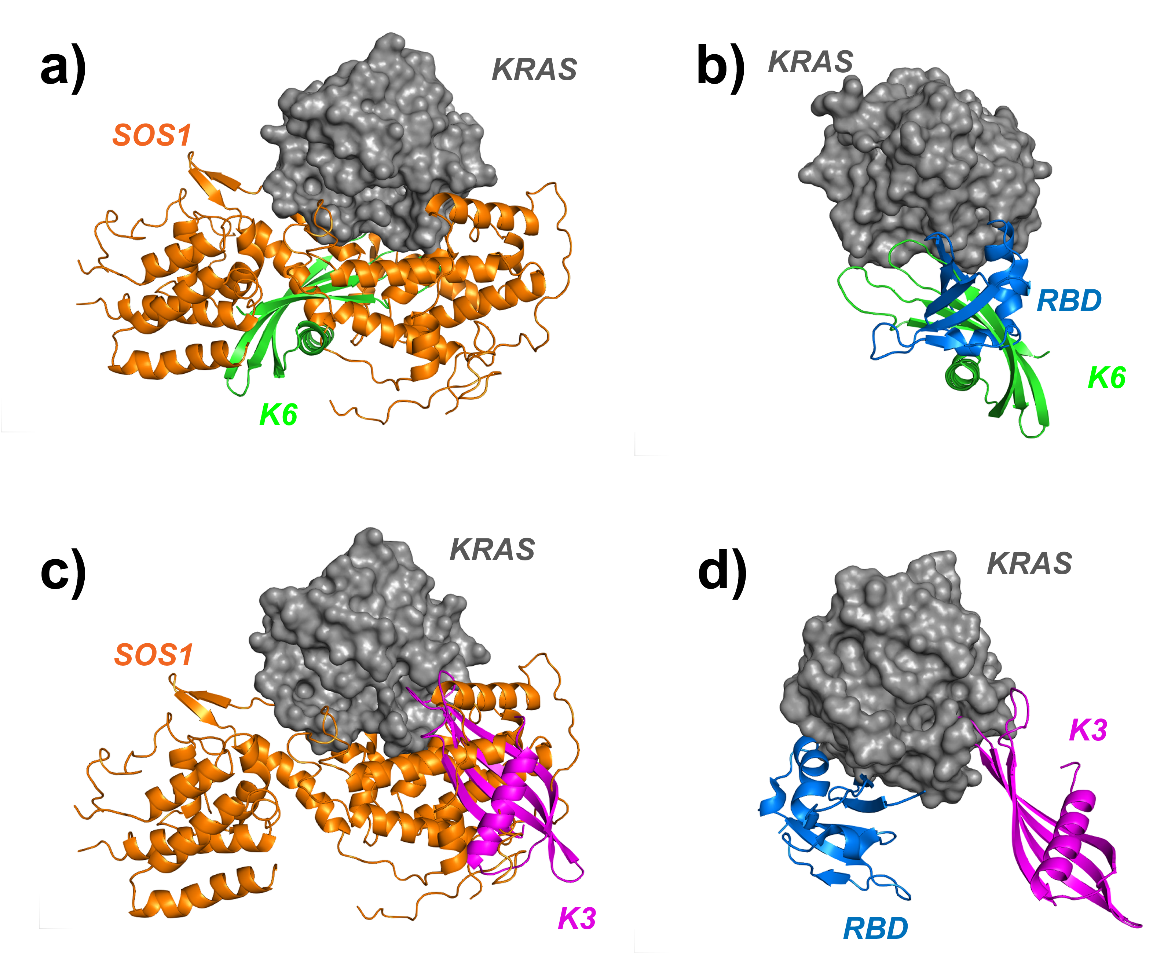

**Supplementary Figure 2. Affimer binding locations compared to other RAS binding biologics.** Affimer K6 is shown in green, Affimer K3 in magenta and RAS in grey. **a)** Compared to NS1 monobody (red) that binds the α4-α5 interface. **b)** DARPins K13 and K19 (yellow and wheat respectively) that bind α3 and α4. **c)** DARPin K27 that spans across the SI/SII pocket (orange). **d)** Intrabody iDab6 (blue) that also spans across the SI/SII pocket**.** Images were generated in MacPyMOLv1.7.2.3 using PDB codes RAS:NS1-5E95; RAS:DARPin K13-6H46; RAS:DARPin K19-6H47; RAS:DARPin K27-5O2S and RAS:iDab6-2UZI

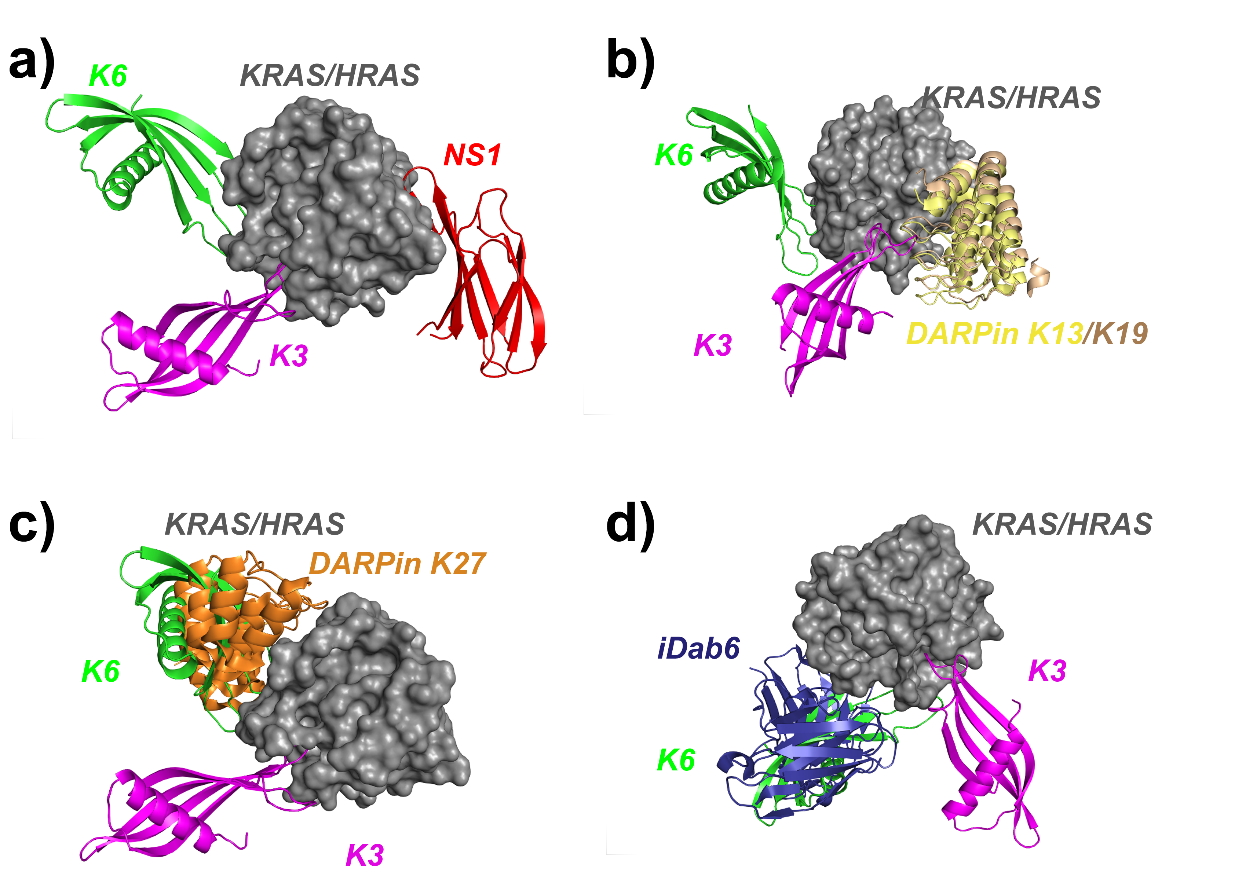

**Supplementary Figure 3. Uncropped images of all membranes shown in this paper.**

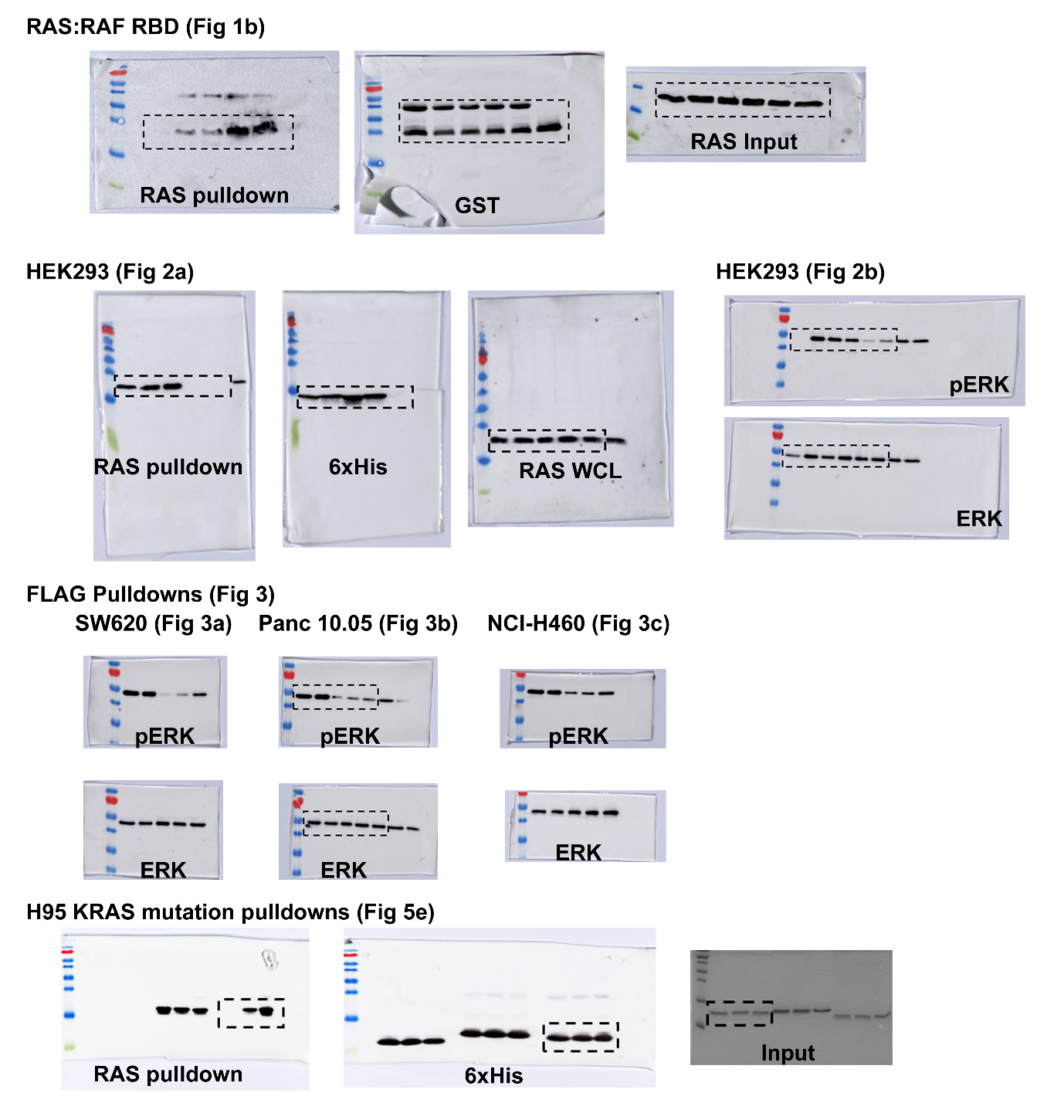

**SUPPLEMENTARY TABLES**

**Supplementary Table 1. Amino acid sequences of variable regions of RAS-binding Affimers**

| **Affimer** | **Variable Region 1** | **Variable Region 2** |
| --- | --- | --- |
| K3 | HSIDIWYDF | KLNNSHTYK |
| K6 | HFTPWFQRN | RIMVTDKMR |
| K19 | FFYLWLAPG | AANSPMYHE |
| K37 | QYNPWFQTN | VIHGTRWGN |
| K68 | VYNPWYQVN | NMRVDMIVH |
| K69 | WHFDYQQYN | RQLRMGSMN |
| K91 | WDFSAWWKY | RNRYFKFPN |

**Supplementary Table 2. Affimer inhibition of SOS1-mediated nucleotide exchange.** IC_50_ values calculated from 3 independent experiments. Data is mean ± SEM.

| **Affimer** | **IC_50_ values (nM)** | | | | | |
| --- | --- | --- | --- | --- | --- | --- |
|  | **KRAS wt** | **KRAS G12D** | **KRAS G12V** | **KRAS Q61H** | **HRAS wt** | **NRAS** |
| K3 | 144 ± 94 | 144 ± 40 | 176 ± 115 | 3005 ± 865 | 2585 ± 335 | No inhibition |
| K6 | 594 ± 271 | 185 ± 46 | 571 ± 148 | 532 ± 165 | 389 ± 187 | 477 ± 44 |
| K37 | 697 ± 158 | 356 ± 161 | 640 ± 253 | 1075 ± 651 | 626 ± 320 | 647 ± 219 |

**Supplementary Table 3. X-ray crystallographic data collection, processing and refinement statistics for Affimer-KRAS complexes.** Values given in parentheses correspond to those in the outermost shell of the resolution range.

§
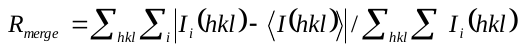

* *R*_p.i.m_ is the precision-indicating (multiplicity-weighted) *R*_merge_ relative to *I*+ or *I*-.

† *R*_free_ was calculated with 5% of the reflections set aside randomly.

ξ Ramachandran analysis using the program Molprobity^38^.

| **Data set** | **K3-KRAS** | **K6-KRAS** |
| --- | --- | --- |
| Source | ESRF ID30A-1 | Diamond Light Source I04-1 |
| Wavelength (Å) | 0.9660 | 0.9159 |
| Resolution range (Å) * | 54.09–2.06 (2.11–2.06) | 64.12-1.90 (1.95-1.90) |
| Space group | *P* 2_1_ | *I* 4*_1_* |
| Unit-cell parameters (Å) | *a*=73.1, *b*=39.5, *c*=113.1 | *a*=*b*=71.6, *c*=144.4 |
|  | α=90.0°,β=106.9°,γ=90.0° | α=90.0°,β=90.0°,γ=90.0° |
| No. of observed reflections | 113731 (8427) | 379142 (24154) |
| No. of unique reflections | 38740 (2830) | 28572 (2116) |
| Redundancy | 2.9 (3.0) | 13.3 (11.4) |
| Completeness (%) * | 99.7 (99.9) | 100.0 (100.0) |
| < *I/σ(I)* >* | 6.2 (1.3) | 13.3 (0.6) |
| *R*_merge_ (%)§* | 8.9 (39.9) | 8.7 (269.0) |
| *R*_pim_ (%) ¥* | 6.3 (29.1) | 2.5 (83.4) |
| CC (1/2) | 0.978 (0.961) | 1.00 (0.799) |
| Resolution range for refinement (Å) | 54.16-2.06 | 64.12-1.90 |
| *R* factor (%) | 23.65 | 21.09 |
| *R*_free_ (%)† | 27.78 | 24.65 |
| No. of protein non-H atoms | 4137 | 2012 |
| No. of water molecules | 103 | 124 |
| R.m.s.d bond lengths (Å) ξ | 0.0055 | 0.0086 |
| R.m.s.d bond angles (˚) ξ | 1.0202 | 1.2904 |
| Average overall *B* factor (Å^2^) |  |  |
| Protein | 46.6 | 67.3 |
| Ligand | 37.5 | 71.4 |
| Water | 45.4 | 68.2 |
| Residues in the favoured region of Ramachandran plot (%) ‡ |  |  |
| Favoured region | 94.8 | 96.8 |
| Outliers | 2 | 0 |
| PDB code | 6YXW | 6YR8 |

**Supplementary Table 4 Primers used in this study**

| **Primer name** | **Primer sequence 5’ – 3’** |
| --- | --- |
| KRAS - forward | CGCGCTAGCATGACCGAATATAAACTGGTGG |
| KRAS - reverse | CGTTGGCGGCCGCTTATTTATGTTTGCGAATTTCACG |
| Affimer-His-forward | ATGGATCCGCCACCATGGCCGCTACCGGTGTTCGTG |
| Affimer-His-reverse | GTTTGGCCAACCACTGCGACTAATATTTCTACTGCTACTGTTCGTAGTAGTAGTAGTAGTAATCCCATTCGCCGGCGATTACG |
| Affimer-GFP-forward | TATATGCGATCGCCATGGGTAACGAAAACTCCCTG |
| Affimer GFP-reverse | AATACGCGTAGCGTCACCAACCGGTTTG |
| K3-VR1.1 forward | GTTGTTAAAGCGAAAGAACAGGCTTCTATCGACATCTGGTACGAC |
| K3-VR1.1 reverse | GTCGTACCAGATGTCGATAGAAGCCTGTTCTTTCGCTTTAACAAC |
| K3-VR1.2 forward | AAAGCGAAAGAACAGCATGCTATCGACATCTGGTACG |
| K3-VR1.2 reverse | CGTACCAGATGTCGATAGCATGCTGTTCTTTCGCTTT |
| K3-VR1.3 forward | GTTAAAGCGAAAGAACAGCATTCTGCTGACATCTGGTACGACTTCACCATG |
| K3-VR1.3 reverse | CATGGTGAAGTCGTACCAGATGTCAGCAGAATGCTGTTCTTTCGCTTTAAC |
| K3-VR1.4 forward | CGAAAGAACAGCATTCTATCGCTATCTGGTACGACTTCACCAT |
| K3-VR1.4 reverse | ATGGTGAAGTCGTACCAGATAGCGATAGAATGCTGTTCTTTCG |
| K3-VR1.5 forward | CGAAAGAACAGCATTCTATCGACGCTTGGTACGACTTCACCATGTACTA |
| K3-VR1.5 reverse | TAGTACATGGTGAAGTCGTACCAAGCGTCGATAGAATGCTGTTCTTTCG |
| K3-VR1.6 forward | AGAACAGCATTCTATCGACATCGCTTACGACTTCACCATGTACTACC |
| K3-VR1.6 reverse | GGTAGTACATGGTGAAGTCGTAAGCGATGTCGATAGAATGCTGTTCT |
| K3-VR1.7 forward | CAGCATTCTATCGACATCTGGGCTGACTTCACCATGTACTACCTG |
| K3-VR1.7 reverse | CAGGTAGTACATGGTGAAGTCAGCCCAGATGTCGATAGAATGCTG |
| K3-VR1.8 forward | ATTCTATCGACATCTGGTACGCTTTCACCATGTACTACCTGAC |
| K3-VR1.8 reverse | GTCAGGTAGTACATGGTGAAAGCGTACCAGATGTCGATAGAAT |
| K3-VR1.9 forward | TCTATCGACATCTGGTACGACGCTACCATGTACTACCTGACCCTG |
| K3-VR1.9 reverse | CAGGGTCAGGTAGTACATGGTAGCGTCGTACCAGATGTCGATAGA |
| K3-VR2.1 forward | CTGTACGAAGCGAAAGTTTGGGTTAAGGCTCTGAACAACAGTCATACCTATAAAAAC |
| K3-VR2.1 reverse | GTTTTTATAGGTATGACTGTTGTTCAGAGCCTTAACCCAAACTTTCGCTTCGTACAG |
| K3-VR2.2 forward | GTACGAAGCGAAAGTTTGGGTTAAGAAAGCTAACAACAGTCATACCTATAAAAACTTC |
| K3-VR2.2 reverse | GAAGTTTTTATAGGTATGACTGTTGTTAGCTTTCTTAACCCAAACTTTCGCTTCGTAC |
| K3-VR2.3 forward | CGAAGCGAAAGTTTGGGTTAAGAAACTGGCTAACAGTCATACCTATAAAAACTTCAAAG |
| K3-VR2.3 reverse | CTTTGAAGTTTTTATAGGTATGACTGTTAGCCAGTTTCTTAACCCAAACTTTCGCTTCG |
| K3-VR2.4 forward | AGCGAAAGTTTGGGTTAAGAAACTGAACGCTAGTCATACCTATAAAAACTTCAAAGAAC |
| K3-VR2.4 reverse | GTTCTTTGAAGTTTTTATAGGTATGACTAGCGTTCAGTTTCTTAACCCAAACTTTCGCT |
| K3-VR2.5 forward | AGTTTGGGTTAAGAAACTGAACAACGCTCATACCTATAAAAACTTCAAAGAAC |
| K3-VR2.5 reverse | GTTCTTTGAAGTTTTTATAGGTATGAGCGTTGTTCAGTTTCTTAACCCAAACT |
| K3-VR2.6 forward | CGAAAGTTTGGGTTAAGAAACTGAACAACAGTGCTACCTATAAAAACTTCAAAG |
| K3-VR2.6 reverse | CTTTGAAGTTTTTATAGGTAGCACTGTTGTTCAGTTTCTTAACCCAAACTTTCG |
| K3-VR2.7 forward | GGTTAAGAAACTGAACAACAGTCATGCTTATAAAAACTTCAAAGAACTGCAGG |
| K3-VR2.7 reverse | CCTGCAGTTCTTTGAAGTTTTTATAAGCATGACTGTTGTTCAGTTTCTTAACC |
| K3-VR2.8 forward | AAGAAACTGAACAACAGTCATACCGCTAAAAACTTCAAAGAACTGCAGGAG |
| K3-VR2.8 reverse | CTCCTGCAGTTCTTTGAAGTTTTTAGCGGTATGACTGTTGTTCAGTTTCTT |
| K3-VR2.9 forward | AAGAAACTGAACAACAGTCATACCTATGCTAACTTCAAAGAACTGCAGGAGTTCAA |
| K3-VR2.9 reverse | TTGAACTCCTGCAGTTCTTTGAAGTTAGCATAGGTATGACTGTTGTTCAGTTTCTT |
| K6-VR1.1 forward | TGTTAAAGCGAAAGAACAGGCTTTCACTCCGTGGTTCCAG |
| K6-VR1.1 reverse | CTGGAACCACGGAGTGAAAGCCTGTTCTTTCGCTTTAACA |
| K6-VR1.2 forward | TCGTGTTGTTAAAGCGAAAGAACAGCATGCTATCCGTGGTTCCAG |
| K6-VR1.2 reverse | CTGGAACCACGGAGTAGCATGCTGTTCTTTCGCTTTAACAACACGA |
| K6-VR1.3 forward | GAAAGAACAGCATTTCGCTCCGTGGTTCCAGCG |
| K6-VR1.3 reverse | CGCTGGAACCACGGAGCGAAATGCTGTTCTTTC |
| K6-VR1.4 forward | TAAAGCGAAAGAACAGCATTTCACTGCTTGGTTCCAGCGTA |
| K6-VR1.4 reverse | TACGCTGGAACCAAGCAGTGAAATGCTGTTCTTTCGCTTTA |
| K6-VR1.5 forward | AAAGAACAGCATTTCACTCCGGCTTTCCAGCGTAACACCATGTAC |
| K6-VR1.5 reverse | GTACATGGTGTTACGCTGGAAAGCCGGAGTGAAATGCTGTTCTTT |
| K6-VR1.6 forward | GAACAGCATTTCACTCCGTGGGCTCAGCGTAACACCATGTACTAC |
| K6-VR1.6 reverse | GTAGTACATGGTGTTACGCTGAGCCCACGGAGTGAAATGCTGTTC |
| K6-VR1.7 forward | CAGCATTTCACTCCGTGGTTCGCTCGTAACACCATGTACTACCTG |
| K6-VR1.7 reverse | CAGGTAGTACATGGTGTTACGAGCGAACCACGGAGTGAAATGCTG |
| K6-VR1.8 forward | CACTCCGTGGTTCCAGGCTAACACCATGTACTACC |
| K6-VR1.8reverse | GGTAGTACATGGTGTTAGCCTGGAACCACGGAGTG |
| K6-VR1.9 forward | CATTTCACTCCGTGGTTCCAGCGTGCTACCATGTACTACCTGACC |
| K6-VR1.9 reverse | GGTCAGGTAGTACATGGTAGCACGCTGGAACCACGGAGTGAAATG |
| K6-VR2.1 forward | CTGTACGAAGCGAAAGTTTGGGTTAAGGCTATTATGGTTACCGATAAAATGAGAAAC |
| K6-VR2.1 reverse | GTTTCTCATTTTATCGGTAACCATAATAGCCTTAACCCAAACTTTCGCTTCGTACAG |
| K6-VR2.2 forward | GAAGCGAAAGTTTGGGTTAAGTGTGCTATGGTTACCGATAAAATGAGAAAC |
| K6-VR2.2 reverse | GTTTCTCATTTTATCGGTAACCATAGCTCTCTTAACCCAAACTTTCGCTTC |
| K6-VR2.3 forward | TGTACGAAGCGAAAGTTTGGGTTAAGAGAATTGCTGTTACCGATAAAATGAGAAACT |
| K6-VR2.3 reverse | AGTTTCTCATTTTATCGGTAACAGACCTTCTCTTAACCCAAACTTTCGCTTCGTACA |
| K6-VR2.4 forward | AAAGTTTGGGTTAAGTGTTAATAGGCTACCGATAAAATGAGAAACTTC |
| K6-VR2.4 reverse | GAAGTTTCTCATTTTATCGGTAGCCATAATTCTCTTAACCCAAACTTT |
| K6-VR2.5 forward | AAAGTTTGGGTTAAGAGAATTATGGTTGCTGATAAAATGAGAAACTTCAAAGAACTG |
| K6-VR2.5 reverse | CAGTTCTTTGAAGTTTCTCATTTTATCAGCAACCATAATTCTCTTAACCCAAACTTT |
| K6-VR2.6 forward | GGTTAAGAGAATTATGGTTACCGCTAAAATGAGAAACTTAAAGAAC |
| K6-VR2.6 reverse | GTTCTTTGAAGTTTCTCATTTTAGCGGTAACCATAATTCTCTTAACC |
| K6-VR2.7 forward | GGGTTAAGAGAATTATGGTTACCGATGCTATGAGAAACTTCAAAGAACTGCAGG |
| K6-VR2.7 reverse | CCTGCAGTTCTTTGAAGTTTCTCATAGCATCGGTAACCATAATTCTCTTAACCC |
| K6-VR2.8 forward | GAAAGTTTGGGTTAAGTGAATTATGGTTACCGATAAAGCTAGAAACTTCAAAGAACTGC |
| K6-VR2.8reverse | GCAGTTCTTTGAAGTTTCTAGCTTTATCGGTAACCATAATTCTCTTAACCCAAACTTTC |
| K6-VR2.9 forward | AAGAGAATTATGGTTACCGATAAAATGGCTAACTTCAAAGAACTGCAGGAGTTCAAA |
| K6-VR2.9 reverse | TTTGAACTCCTGCAGTTATTTGAAGTTAGCCATTTTATCGGTAACCATAATTCTCTT |
| Affimer-VR1 forward | ATGGCTAGCAACTCCCTGGAAATCGAAG |
| Affimer-VR1 reverse | CACCGTCTTTAGCTTCCAGG |
| Affimer-VR2 forward | CCTGGAAGCTAAAGACGGTG |
| Affimer-VR2 reverse | TACCCTAGTGGTGATGATGGTGATGC |
